# Identification of Viral Activators of the HSV-2 UL13 Protein Kinase

**DOI:** 10.1101/2025.07.06.663391

**Authors:** Naoto Koyanagi, Kosuke Takeshima, Saori Shio, Yuhei Maruzuru, Akihisa Kato, Yasushi Kawaguchi

## Abstract

Although previous studies reported that the herpes simplex virus 2 (HSV-2) UL13 protein kinase mediates the phosphorylation of elongation factor 1δ (EF-1δ) in infected cells, we show here that individual expression of UL13 was insufficient to induce phosphorylation of EF-1δ in mammalian cells. This led us to hypothesize that HSV-2 UL13 requires viral cofactors for full kinase activity and prompted us to identify such cofactors. Our results were as follows. (i) Co-expression of UL13 with UL55 or Us10 significantly enhanced phosphorylation of EF-1δ compared to UL13 alone. (ii) UL13 was co-precipitated with UL55 or Us10 upon co-expression, and its kinase activity was significantly increased in their presence, as demonstrated by *in vitro* kinase assays. (iii) In HSV-2-infected cells, UL13 was specifically co-precipitated with Us10 and UL55. (iv) The UL55-null mutation significantly reduced phosphorylation of EF-1δ in HSV-2-infected cells, whereas the Us10-null mutation had little effect; however, the double-null mutation further decreased the phosphorylation compared to the UL55-null mutation alone. (v) The UL55-null mutation, but not the Us10-null mutation, significantly reduced HSV-2 replication and cell-cell spread in U2OS cells to levels comparable to those observed with the UL13 kinase-dead mutation. These results suggest that UL55 acts as a principal activator of UL13 in HSV-2-infected cells, whereas Us10 serves as an auxiliary activator. Moreover, the role of UL13 kinase activity in HSV-2 replication and cell-cell spread in U2OS cells appears to be largely dependent on UL55.

**Importance:** Herpesviruses encode conserved protein kinases (CHPKs) that often target cellular cyclin-dependent kinase (CDK) phosphorylation sites. CHPKs from beta-and gammaherpesviruses can exhibit these CDK-like functions even when individually expressed in mammalian cells. In contrast, CHPKs from alphaherpesviruses display these CDK-like functions in infected cells, but not upon individual expression, suggesting that they require additional viral factors to exhibit full kinase activity. In this study, we focused on HSV-2 UL13, an alphaherpesvirus CHPK, and identified HSV-2 UL55 and Us10 as viral activators of UL13. In HSV-2-infected cells, UL55 functions as a principal activator of UL13, while Us10 serves as an auxiliary activator. Importantly, the contribution of UL13 kinase activity to HSV-2 replication and cell-cell spread appears to be largely dependent on the presence of UL55. Our findings uncover a previously unrecognized mechanism of CHPK regulation in alphaherpesviruses and provide new insights into the evolutionary diversification of viral kinase control.

## Introduction

Viruses in the family *Herpesviridae* (herpesviruses) are subclassified into three subfamilies, *Alphaherpesvirinae*, *Betaherpesvirinae,* and *Gammaherpesvirinae*, based on molecular and biological properties (1). Although members of these subfamilies exhibit a wide range of pathogenicity, clinical manifestations, and biological characteristics (1), their genomes encode a number of conserved viral proteins (1). This conservation suggests that these viral proteins play fundamental and universal roles in the life cycles of herpesviruses. Herpes simplex virus 2 (HSV-2), a member of the subfamily *Alphaherpesvirinae*, encodes UL13 protein kinase, a serine/threonine protein kinase that is one of the conserved viral proteins throughout the *Herpesviridae* family (2, 3). These conserved viral protein kinases, designated conserved herpesvirus protein kinases (CHPKs) (2, 3), include UL13 of HSV-1 and open reading frame 47 (ORF47) of varicella-zoster virus (VZV) in the *Alphaherpesvirinae* subfamily; UL97 of human cytomegalovirus (HCMV), and U69 of human herpesvirus 6A (HHV-6A) and 6B (HHV-6B), and 7 (HHV-7) in the *Betaherpesvirinae* subfamily; and BGLF4 of Epstein-Barr virus (EBV) and ORF36 of Kaposi’s sarcoma-associated herpesvirus (KSHV) in the *Gammaherpesvirinae* subfamily. CHPKs have been implicated in various aspects of viral replication and pathogenicity, underscoring their importance in the life cycles of herpesviruses (2, 3).

CHPKs have been reported to share functional and regulatory similarities with cellular cyclin-dependent kinases (CDKs) and are therefore also referred to as viral CDK-like kinases (2–12). Accumulating evidence suggests that CHPKs target a number of CDK phosphorylation sites (4–9, 12–15) and that phosphorylation of conserved tyrosine residues within the GxGxxG motifs of representative CHPKs from the three subfamilies of the *Herpesviridae* family negatively regulates their kinase activities (16) similar to the regulatory phosphorylation observed in CDKs (17). However, previous studies on the CDK-like functions of CHPKs have led to conflicting interpretations, particularly for CHPKs of alphaherpesviruses. Thus, it has been reported that, in HSV-1 or HSV-2-infected cells, UL13 phosphorylates CDK1 phosphorylation site (Ser-133) of cellular translation elongation factor 1δ (EF-1δ) (4, 18). The ability to phosphorylate EF-1δ Ser-133 was also demonstrated for HCMV UL97 and EBV BGLF4, based on the observations that individual expression of these viral protein kinases in mammalian cells induces phosphorylation of this site (19, 20). These findings suggest that the CDK-like functions of CHPKs are conserved among CHPKs from all subfamilies of the *Herpesviridae* family. In contrast, the ability to phosphorylate CDK target sites in cellular retinoblastoma protein (Rb) and sterile alpha motif and HD domain 1 (SAMHD1) upon individual CHPK expression in mammalian cells is shared by CHPKs from beta-and gammaherpesviruses, but not by CHPKs from alphaherpesviruses (12, 21). It remains unclear whether these discrepancies arise from differences in CHPK substrate specificity or from variations in experimental systems, such as CHPK expression in the context of viral infection versus transient expression in mammalian cells.

In this study, we sought to address these discrepancies. Whereas phosphorylation of EF-1δ at Ser-133 in HSV-2-infected cells has been reported (18), we demonstrated here that individual expression of HSV-2 UL13 in mammalian cells was insufficient to induce phosphorylation of EF-1δ at Ser-133, similar to what was previously observed for Rb and SAMHD1, which were also not phosphorylated upon individual expression of CHPKs from alphaherpesviruses in mammalian cells (12, 21). These findings led us to hypothesize that HSV-2 UL13 may require one or more viral cofactors to exhibit full kinase activity. To identify such cofactors, we screened HSV-2 tegument proteins and found that UL55 and Us10 act as viral activators of HSV-2 UL13.

## Results

### Individual expression of HSV-2 UL13 and VZV ORF47 in COS-7 cells was insufficient to induce phosphorylation of EF-1δ at Ser-133

To examine whether individual expression of CHPKs in mammalian cells can induce phosphorylation of EF-1δ at Ser-133, simian kidney epithelial COS-7 cells were transfected with a plasmid expressing Flag-tagged EF-1δ fused to enhanced green fluorescent protein (EGFP) [EGFP-EF-1δ(F)] (18) in combination with each of the plasmids expressing wild-type CHPKs tagged with Strep-tag (SE-CHPKs) (16), and then subjected to immunoblotting with a monoclonal antibody specific for EF-1δ phosphorylated at Ser-133 (EF-1δ-S133^P^) (18). Phosphorylation of EGFP-EF-1δ(F) at Ser-133 was increased upon individual expression of SE-CHPKs from beta-and gammaherpesviruses, including HCMV SE-UL97, HHV-6B SE-U69, EBV SE-BGLF4 and KSHV SE-ORF36, but not from alphaherpesviruses, namely HSV-2 SE-UL13 and VZV SE-ORF47 (Fig. 1A). Notably, the expression level of HSV-2 SE-UL13 appeared higher than those of HCMV SE-UL97, EBV SE-BGLF4 and KSHV SE-ORF36, and the expression level of VZV SE-ORF47 was comparable to that of HCMV SE-UL97 (Fig. 1A). Nevertheless, these alphaherpesvirus SE-CHPKs were insufficient to induce phosphorylation of EF-1δ at Ser-133 upon individual expression. These results are in agreement with earlier observations that CHPKs from beta-and gammaherpesviruses, but not from alphaherpesviruses, can induce phosphorylation of CDK target sites in Rb and SAMHD1 when individually expressed in mammalian cells (12, 21). These results, together with earlier observations that HSV-2 UL13 can induce phosphorylation of EF-1δ at Ser-133 in infected cells (18), led us to hypothesize that UL13 may require a viral cofactor(s) to exhibit full kinase activity and prompted us to identify such cofactors.

**Fig. 1.**
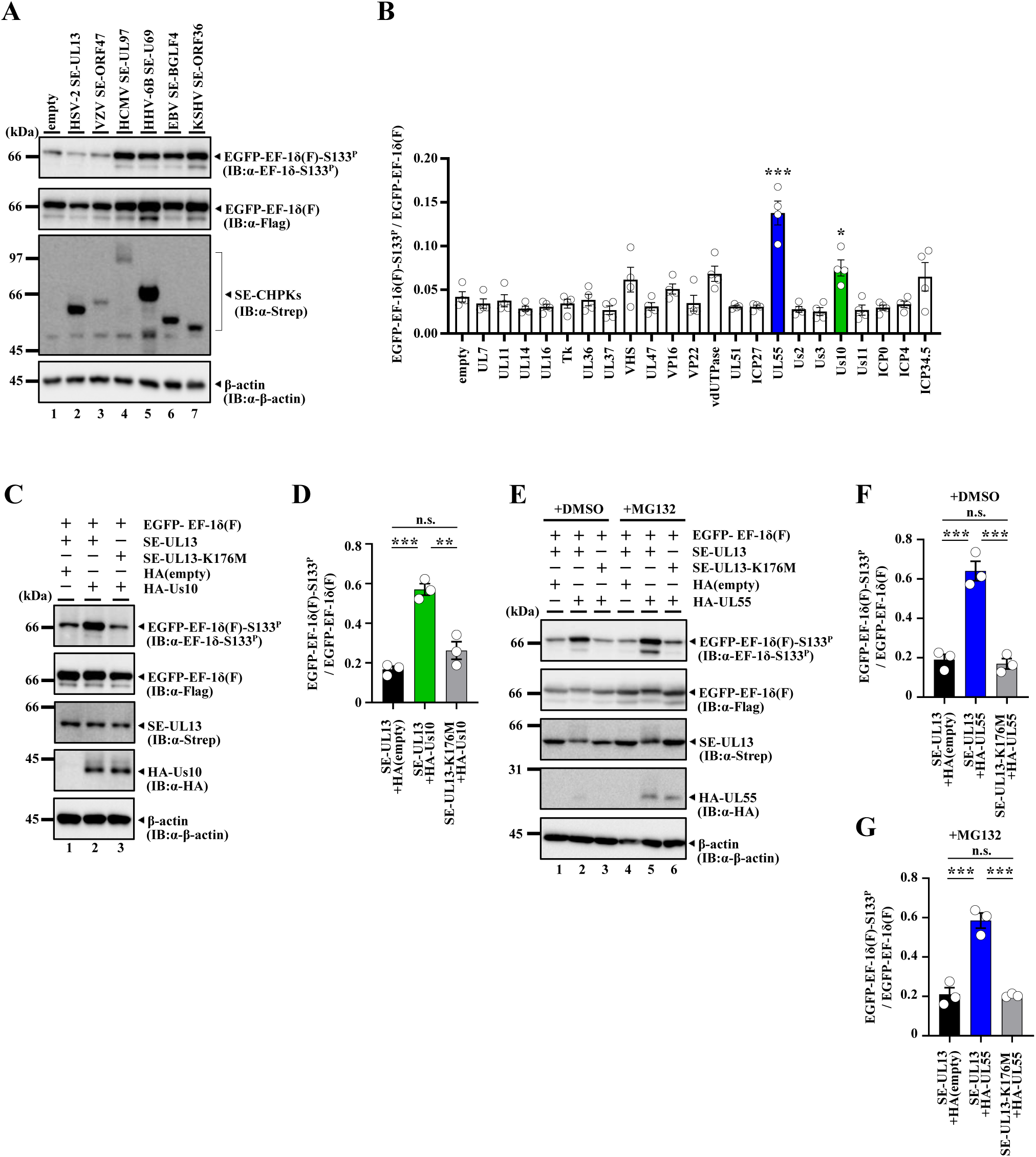
Identification of HSV-2 UL55 and Us10 as an activator of HSV-2 UL13 kinase activity. **A.** COS-7 cells were transfected with a plasmid expressing EGFP-EF-1δ(F) (lanes 1-7) in combination with an empty plasmid (lane 1), SE-UL13 (lanes 2), SE-ORF47 (lane 3), SE-UL97 (lane 4), SE-U69 (lane 5), SE-BGLF4 (lane 6), or SE-ORF36 (lane 7), and harvested 48 h post-transfection. Cell lysates were analyzed by immunoblotting with antibodies to Flag-tag, EF-1δ-S133^P^, Strep-tag, or β-actin. Digital images are representative of three independent experiments. **B.** COS-7 cells were transfected with plasmids expressing EGFP-EF-1δ(F), SE-UL13 and one of the 22 HSV-2 proteins or an empty plasmid, and harvested 48 h post-transfection. Cell lysates were analyzed by immunoblotting with antibodies to Flag-tag or EF-1δ-S133^P^. Amount of EGFP-EF-1δ(F)-S133^P^ protein detected with anti-EF-1δ-S133^P^ monoclonal antibody relative to that of EGFP-EF-1δ(F) protein detected with anti-Flag antibody in transfected cells. Data were normalized by dividing the sum of the data on the same blot (52). Each value is the mean ± SEM of four experiments. Statistical significance was analyzed by one-way ANOVA with Dunnett’s multiple comparisons test comparing to the empty plasmid. Asterisks indicate statistically significant values (*, *P* < 0.05; ***, *P* < 0.001). **C.** COS-7 cells were transfected with a plasmid expressing EGFP-EF-1δ(F) in combination with a plasmid expressing SE-UL13 (lanes 1, 2) or SE-UL13-K176M (lane 3), and an empty plasmid (lane 1) or a plasmid expressing HA-Us10 (lanes 2, 3), harvested 48 h post-transfection, and lysates were then analyzed by immunoblotting with the indicated antibodies. Digital images are representative of three independent experiments. **D.** Amount of EGFP-EF-1δ(F)-S133^P^ protein detected with anti-EF-1δ-S133^P^ monoclonal antibody (C, top panel) relative to that of EGFP-EF-1δ(F) protein detected with anti-Flag antibody (C, second panel from the top) in transfected cells. Data were normalized by dividing the sum of the data on the same blot (52). Each value is the mean ± SEM of three experiments. Statistical significance was analyzed by ANOVA with Tukey’s test. Asterisks indicate statistically significant values (**, *P* < 0.01, ***, *P* < 0.001). n.s., not significant. **E.** COS-7 cells were transfected with a plasmid expressing EGFP-EF-1δ(F) in combination with a plasmid expressing SE-UL13 (lanes 1, 2, 4, 5) or SE-UL13-K176M (lanes 3, 6), and an empty plasmid (lane 1 or 4) or a plasmid expressing HA-UL55 (lanes 2, 3, 5, 6). Transfected cells were incubated with DMSO or 10 μM MG132 24 h post-transfection, harvested 48 h post-transfection, and lysates were then analyzed by immunoblotting with the indicated antibodies. Digital images are representative of three independent experiments. **F, G**. Amount of EGFP-EF-1δ(F)-S133^P^ protein detected with anti-EF-1δ-S133^P^ monoclonal antibody (E, top panel) relative to that of EGFP-EF-1δ(F) protein detected with anti-Flag antibody (E, second panel from the top) in transfected cells with DMSO (F) or MG132 (G). Data were normalized by dividing the sum of the data on the same blot (52). Each value is the mean ± SEM of three experiments. Statistical significance was analyzed by ANOVA with Tukey’s test. Asterisks indicate statistically significant values (***, *P* < 0.001). n.s., not significant.

### Identification of viral activators of HSV-2 UL13

CHPKs have been reported to be packaged into the tegument of virions (22–27), an amorphous compartment located between the nucleocapsid and the envelope that contains more than 20 different viral proteins (28). It has been suggested that CHPKs in the tegument are released into the cytoplasm of newly infected cells, where they contribute to establishing conditions favorable for viral replication immediately after viral entry (26, 29). Given this, a viral cofactor(s) for HSV-2 is likely to reside within the tegument compartment. Based on these observations, we focused on HSV-2 tegument proteins and screened them in an attempt to identify viral cofactors that activate UL13.

COS-7 cells were transfected with a plasmid expressing EGFP-EF-1δ(F) together with a plasmid expressing SE-UL13 in combination with each of the plasmids expressing 22 different tegument proteins fused to EGFP, and then subjected to immunoblotting with an anti-EF-1δ-S133^P^ antibody. Among the tegument proteins tested, only UL55 and Us10 significantly enhanced phosphorylation of EGFP-EF-1δ(F) at Ser-133 when co-expressed with SE-UL13 (Fig. 1B).

To verify whether Us10 enhances phosphorylation of EGFP-EF-1δ(F) at Ser-133 mediated by UL13, COS-7 cells were transfected with a plasmid expressing EGFP-EF-1δ(F), together with a plasmid expressing either SE-UL13 or SE-UL13-K176M, a kinase-dead mutant of UL13 (30), in combination with a plasmid expressing HA-tagged Us10 (HA-Us10), and then subjected to immunoblotting with an anti-EF-1δ-S133^P^ antibody. Phosphorylation of EGFP-EF-1δ(F) at Ser-133 was significantly enhanced by co-expression of SE-UL13 with HA-Us10, but not by co-expression of SE-UL13-K176M with HA-Us10 (Fig. 1C and D). To determine whether UL55 similarly enhanced phosphorylation of EGFP-EF-1δ(F) at Ser-133 mediated by UL13, we attempted similar experiments using a plasmid expressing HA-UL55. However, we noted that expression of HA-UL55 was barely detectable. This suggested that expression of HA-UL55 in COS-7 cells may be unstable, possibly due to proteasome-mediated degradation of the viral protein. Therefore, COS-7 cells were transfected with a plasmid expressing EGFP-EF-1δ(F), together with a plasmid expressing SE-UL13 or SE-UL13-K176M, in combination with a plasmid expressing HA-UL55, and treated with either the proteasome inhibitor MG132 or its solvent control DMSO. The cells were then subjected to immunoblotting with an anti-EF-1δ-S133^P^ antibody. Phosphorylation of EGFP-EF-1δ(F) at Ser-133 was significantly enhanced by co-expression of SE-UL13 with HA-UL55, but not by co-expression of SE-UL13-K176M with HA-UL55, regardless of MG132 treatment (Fig. 1E to G). Notably, MG132 treatment led to a marked accumulation of HA-UL55 (Fig. 1E). These results indicate that UL55 and Us10 can enhance the ability of UL13 to phosphorylate EF-1δ at Ser-133. Furthermore, our findings demonstrate that UL55 is inherently unstable when expressed alone in COS-7 cells, likely due to its susceptibility to proteasome-mediated degradation.

To examine whether Us10 or UL55 can activate UL13 kinase activity, human embryonic kidney 293T (HEK293T) cells were transfected with a plasmid expressing SE-UL13 or its kinase-dead mutant SE-UL13-K176M, together with a plasmid expressing Us10 fused to EGFP (Us10-EGFP), UL55 fused to EGFP (UL55-EGFP) or EGFP. Cells were harvested, solubilized, and subjected to pulldown using Strep-Tactin Sepharose beads. The precipitates were incubated in kinase buffer with purified maltose binding protein (MBP) fused to a domain of EF-1δ containing Ser-133 {MBP-EF-1δ(107–146)} or its mutant version MBP-EF-1δ(107–146)-S133A, in which Ser-133 in MBP-EF-1δ(107–146) was substituted with alanine (4), and then analyzed by immunoblotting with an anti-EF-1δ-S133^P^ antibody.

As shown in Fig. 2A, Us10-EGFP was co-precipitated with both SE-UL13 and SE-UL13-K176M. Phosphorylation of MBP-EF-1δ(107–146) at Ser-133, but not that of the S133A mutant, was detected when SE-UL13 was co-expressed with Us10-EGFP. No phosphorylation of MBP-EF-1δ(107–146) at Ser-133 was observed when SE-UL13-K176M was co-expressed with Us10-EGFP or when SE-UL13 was co-expressed with EGFP (Fig. 2A). In contrast to the co-precipitation with Us10-EGFP, UL55-EGFP was co-precipitated with SE-UL13 but not with SE-UL13-K176M (Fig. 2B). Phosphorylation of MBP-EF-1δ(107–146) at Ser-133, but not that of the S133A mutant, was detected when SE-UL13 was co-expressed with UL55-EGFP (Fig. 2B). In contrast, no phosphorylation of MBP-EF-1δ(107–146) at Ser-133 was observed when SE-UL13-K176M was co-expressed with Us10-EGFP or when SE-UL13 was co-expressed with EGFP (Fig. 2B). These results indicate that both Us10 and UL55 can enhance the kinase activity of UL13 and function as viral cofactors that activate UL13, thereby enabling efficient phosphorylation of specific substrates such as EF-1δ. Notably, while SE-UL13 co-expressed with Us10-EGFP was detected as a single band in immunoblotting (Fig. 2A), SE-UL13 co-expressed with UL55-EGFP was detected as two bands with different mobilities in immunoblotting (Fig. 2B): one appeared to correspond to the band observed with Us10-EGFP, and the other exhibited slower mobility. The slower-migrating band of UL13 was previously reported to result from autophosphorylation (18), supporting our conclusion that UL55 activates UL13.

**Fig. 2.**
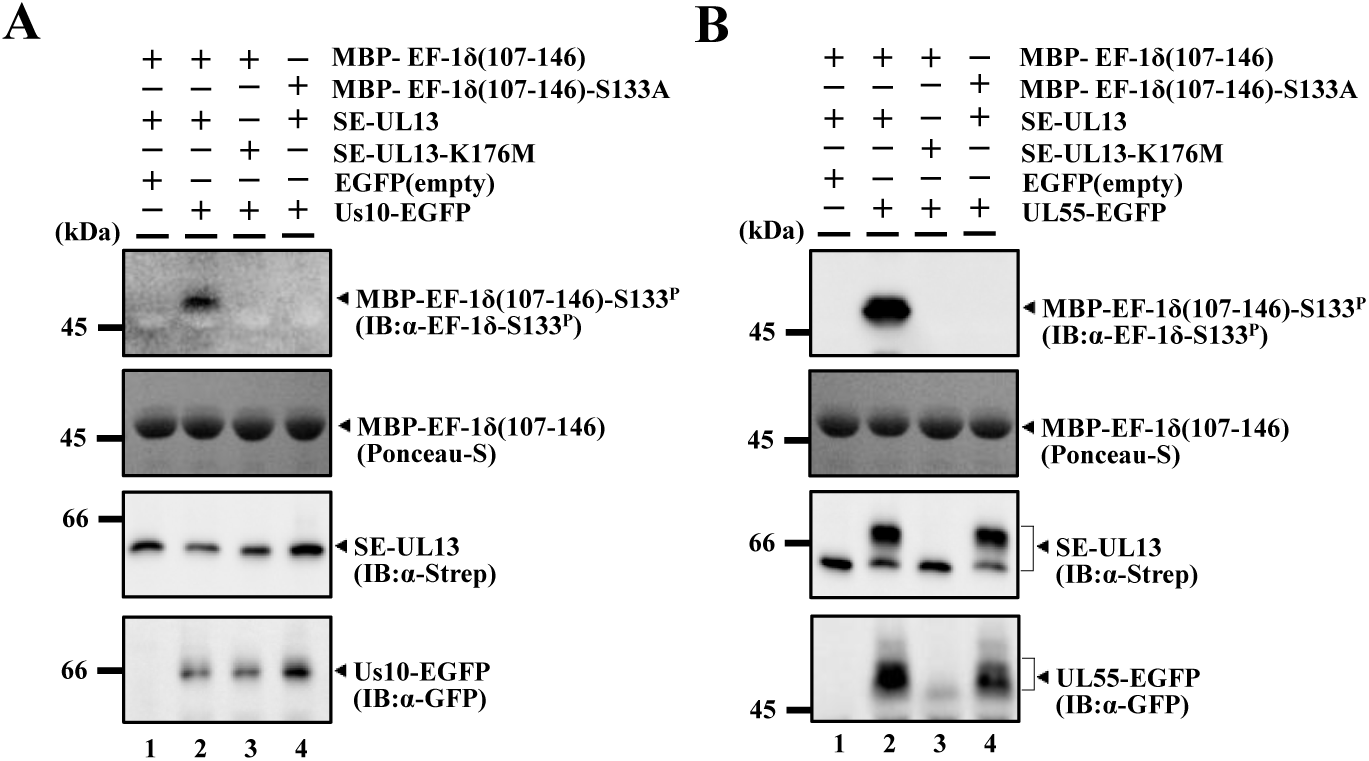
UL55 and Us10 upregulate UL13 kinase activity. **A.** HEK293T cells were transfected with a plasmid expressing SE-UL13 (lanes 1, 2, 4) or SE-UL13-K176M (lane 3), and an empty plasmid (lane 1) or a plasmid expressing Us10-EGFP (lane 2, 3, 4). Transfected cells were harvested 48 h post-transfection, precipitated with StrepTactin-sepharose. For *in vitro* kinase assays, the precipitates were incubated in kinase buffer containing MBP-EF-1δ (107–146) (lane1, 2, 3) or MBP-EF-1δ (107–146)-S133A (lane 4), separated on a denaturing gel, transferred onto a nitrocellulose membrane, subjected to Ponceau-S staining, and then analyzed by immunoblotting with the indicated antibodies. Digital images are representative of three independent experiments. **B.** HEK293T cells were transfected with a plasmid expressing SE-UL13 (lanes 1, 2, 4) or SE-UL13-K176M (lane 3), and an empty plasmid (lane 1) or a plasmid expressing UL55-EGFP (lane 2, 3, 4). Transfected cells were incubated with 10 μM MG132 24 h post-transfection, harvested 48 h post-transfection, precipitated with StrepTactin-sepharos. For *in vitro* kinase assays, the precipitates were incubated in kinase buffer containing MBP-EF-1δ (107–146) (lane1, 2, 3) or MBP-EF-1δ (107–146)-S133A (lane 4), separated on a denaturing gel, transferred onto a nitrocellulose membrane, subjected to Ponceau-S staining, and then analyzed by immunoblotting with the indicated antibodies. Digital images are representative of three independent experiments.

### Construction and characterization of recombinant viruses

To investigate the effects of UL55 and/or Us10 on UL13 in HSV-2-infected cells, we constructed a series of recombinant viruses including the following: a recombinant virus YK873 (UL13-HA) expressing HA-tagged UL13 (UL13-HA); a UL55-null mutant virus YK874 (ΔUL55); a Us10-null mutant virus YK876 (ΔUs10); and a UL55/Us10-double null mutant virus YK878 (ΔUL55/ΔUs10) (Fig. 3). In addition, we generated recombinant viruses in which each of these null mutations was repaired: YK875 (ΔUL55-repair), YK877 (ΔUs10-repair) and YK879 (ΔUL55/ΔUs10-repair) (Fig. 3).

**Fig. 3.**
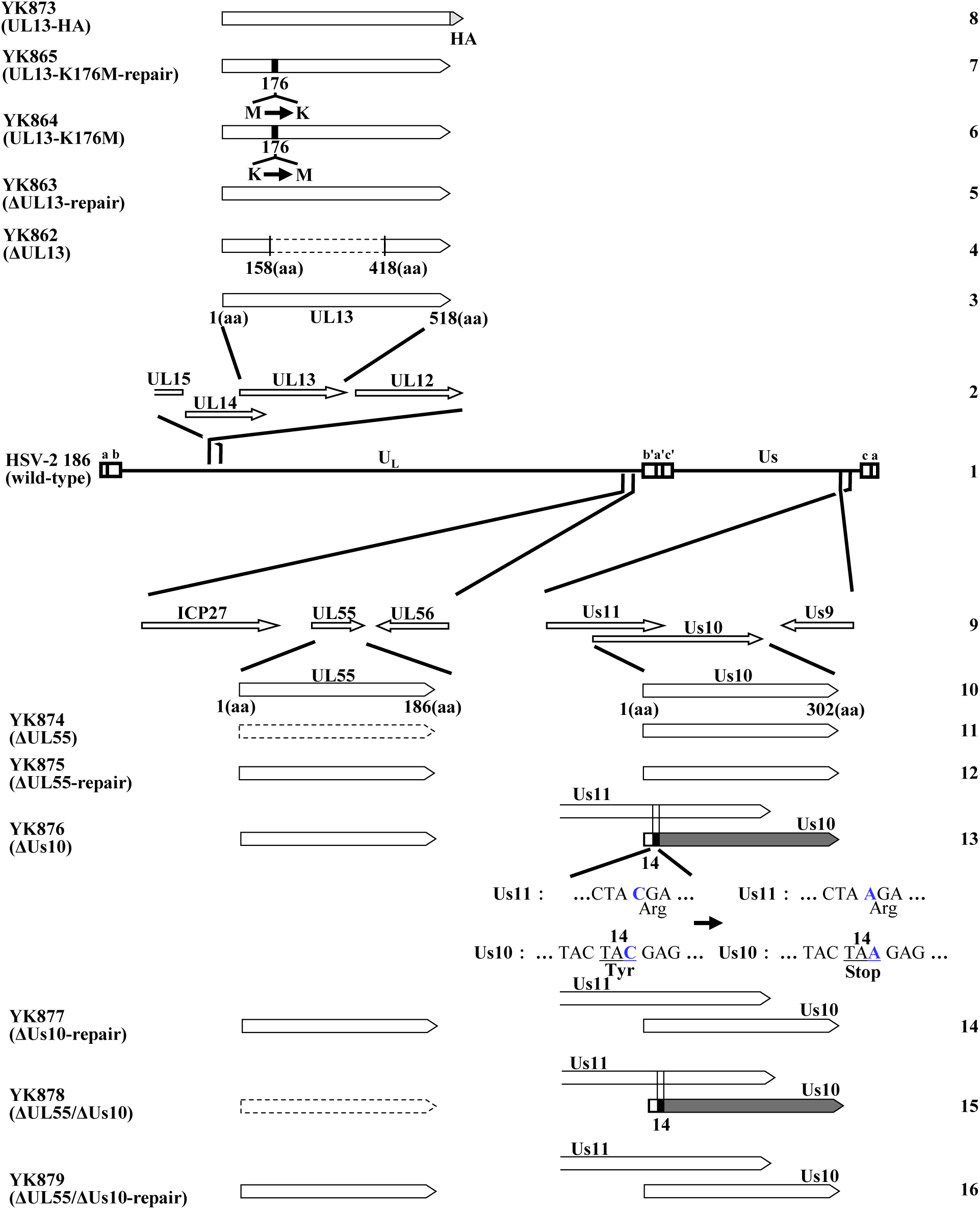
Schematic diagrams of the genome structures of wild-type HSV-2 186 and the relevant domains of the recombinant viruses used in this study. Line 1, wild-type HSV-2 186 genome; Line 2, domain of the UL12 gene to the UL15 gene; Line 3, domain of the UL13 gene; Lines 4 to 8, recombinant viruses with mutations in the UL13 gene; Line 9, domains of UL54 (ICP27) to UL56 and Us9 to Us11 genes; Line 10, domains of the UL55 and Us10 genes; Lines 11 to 16, recombinant viruses with mutations in the UL55 and/or Us10 genes.

The recombinant viruses were characterized as follows. (i) UL13-HA was detected in lysates of simian kidney epithelial Vero cells infected with YK873 (UL13-HA), but not in those infected with wild-type HSV-2 186 (Fig. 4A). (ii) Vero cells infected with wild-type HSV-2 186 or YK875 (ΔUL55-repair) expressed UL55, whereas cells infected with YK874 (ΔUL55) did not (Fig. 4B), confirming that the UL55 gene was successfully disrupted in YK874 (ΔUL55). (iii) Vero cells infected with wild-type HSV-2 186, YK874 (ΔUL55) or YK875 (ΔUL55-repair) produced comparable levels of UL56 and of ICP27, which is encoded by the UL54 gene (Fig. 4C), indicating that the UL55-null mutation had little effect on the expression of its neighboring genes. Notably, UL56 in YK874 (ΔUL55)-infected cells was predominantly detected as a single band in immunoblotting, whereas UL56 in wild-type HSV-2 186-or YK875 (ΔUL55-repair)-infected cells was detected as two bands with different mobilities (Fig. 4C), suggesting that UL55 is required for efficient post-translational modification(s) of UL56 in HSV-2-infected cells. (iv) Vero cells infected with wild-type HSV-2 186 or YK877 (ΔUs10-repair) expressed Us10, whereas cells infected with YK876 (ΔUs10) did not (Fig. 4D), confirming that the Us10 gene was successfully disrupted in YK876 (ΔUs10). (v) Vero cells infected with wild-type HSV-2 186, YK876 (ΔUs10) or YK877 (ΔUs10-repair) produced comparable levels of Us9 and Us11 (Fig. 4E), indicating that the null mutation in YK876 (ΔUs10) has little effect on expression of its neighboring genes. (vi) Vero cells infected with wild-type HSV-2 186 or YK879 (ΔUL55/ΔUs10-repair) expressed both UL55 and Us10, whereas those infected with YK878 (ΔUL55/ΔUs10) did not (Fig. 4F), confirming that both genes were successfully disrupted in YK878 (ΔUL55/ΔUs10). (vii) Vero cells infected with wild-type HSV-2 186, YK878 (ΔUL55/ΔUs10) or YK879 (ΔUL55/ΔUs10-repair) produced comparable levels of ICP27, UL56, Us9 and Us11 (Fig. 4G), indicating that the null mutation in YK878 (ΔUL55/ΔUs10) had little effect on the expression of genes neighboring the UL55 and Us10 loci.

**Fig. 4.**
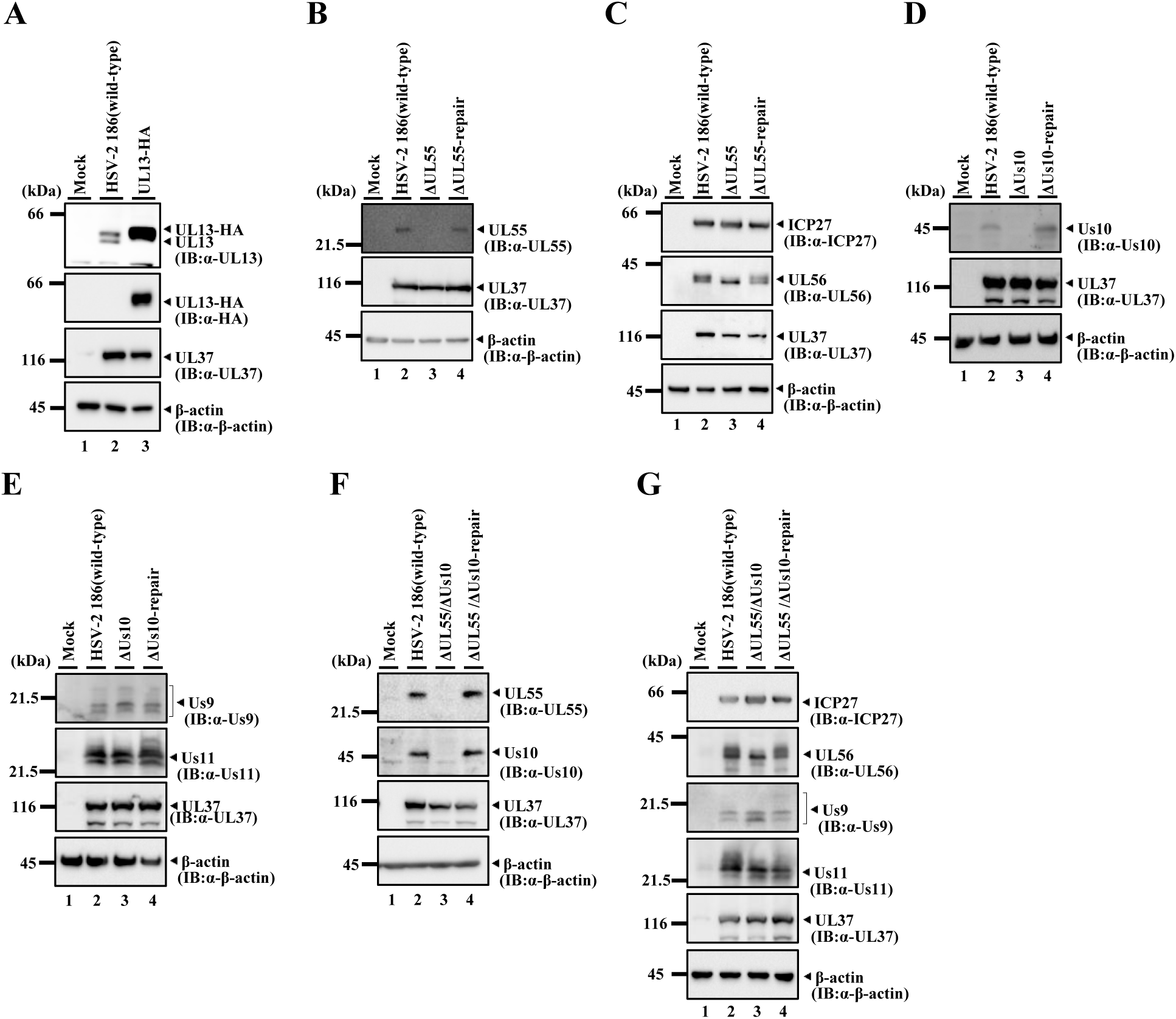
Characterization of the recombinant viruses. **A.** Vero cells were mock infected (lane 1) or infected with wild-type HSV-2 186 (lane 2) or YK873 (UL13-HA) (lane 3) at an MOI of 3 for 24 h and then analyzed by immunoblotting with the indicated antibodies. **B, C.** Vero cells were mock-infected (lane 1) or infected with wild-type HSV-2 186 (lane 2), YK874 (ΔUL55) (lane 3) or YK875 (ΔUL55-repair) (lane 4) at an MOI of 3, harvested at 24 h post-infection, and lysates were analyzed by immunoblotting with the indicated antibodies. **D, E.** Vero cells were mock-infected (lane 1) or infected with wild-type HSV-2 186 (lane 2), YK876 (ΔUs10) (lane 3) or YK877 (ΔUs10-repair) (lane 4) at an MOI of 3, harvested at 24 h post-infection, and lysates were analyzed by immunoblotting with the indicated antibodies. **F, G.** Vero cells were mock-infected (lane 1) or infected with wild-type HSV-2 186 (lane 2), YK878 (ΔUL55/ΔUs10) (lane 3) or YK879 (ΔUL55/ΔUs10-repair) (lane 4) at an MOI of 3, harvested at 24 h post-infection, and lysates were analyzed by immunoblotting with the indicated antibodies. Digital images are representative of three independent experiments.

### Association of UL13 with UL55 and Us10 in HSV-2-infected cells

To investigate whether UL13 interacts with UL55 or Us10 in HSV-2-infected cells, Vero cells were infected with wild-type HSV-2 186 or YK873 (UL13-HA), lysed and immunoprecipitated with an anti-HA antibody. The immunoprecipitates were then analyzed by immunoblotting. As shown in Fig. 5, the anti-HA antibody co-precipitated UL55 and Us10 with UL13-HA from lysates of YK873 (UL13-HA)-infected cells but did not co-precipitate the capsid protein VP23. No such co-precipitation was observed in lysates of wild-type HSV-2 186-infected cells (Fig. 5). These results indicate that UL13 specifically interacts with UL55 and Us10 in HSV-2-infected cells and are in agreement with our findings above that UL55 and Us10 were co-precipitated with UL13 when co-expressed individually with UL13.

**Fig. 5.**
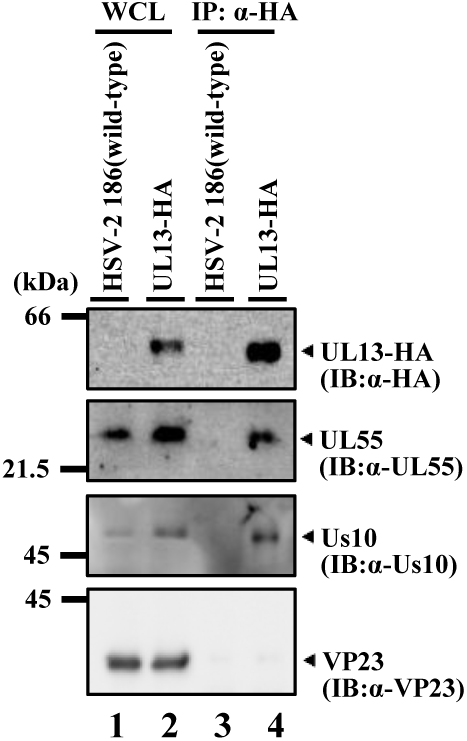
Interactions of UL13 with UL55 and Us10 in HSV-2-infected cells. Vero cells were infected with wild-type HSV-2 186 (lane 1, 3) or YK873 (UL13-HA) (lane 2, 4) at an MOI of 3 for 24 h, lysed, immunoprecipitated with anti-HA antibody, and analyzed by immunoblotting with the indicated antibodies. WCL, whole-cell lysate. Digital images are representative of three independent experiments.

### Effects of UL13 on accumulation of UL55 and Us10 in HSV-2-infected cells

To examine the effects of UL13 on UL55 and Us10 in HSV-2-infected cells, Vero cells were mock-infected or infected with wild-type HSV-2 186, a UL13-null mutant virus YK862 (ΔUL13) (18), its repaired virus YK863 (ΔUL13-repair), YK864 (UL13-K176M) encoding an enzymatically inactive mutant of UL13 (18) or its repaired virus YK865 (UL13-K176M-repair), lysed and subjected to immunoblotting. As shown in Fig. 6A, UL55 and Us10 were barely detectable in cells infected with YK862 (ΔUL13), although they accumulated in cells infected with wild-type HSV-2 186 or YK863 (ΔUL13-repair). Similarly, Us10 was barely detectable in cells infected with YK864 (UL13-K176M), although it accumulated in cells infected with wild-type HSV-2 186 or YK865 (UL13-K176M-repair) (Fig. 6B). In contrast, UL55 accumulated in cells infected with YK864 (UL13-K176M) at levels comparable to those in cells infected with wild-type HSV-2 186 or YK865 (UL13-K176M-repair) (Fig. 6B). As previously reported (18) and shown in Fig. 9A below, the K176M mutation in UL13 does not decrease the accumulation level of UL13 in HSV-2-infected Vero cells. These results indicate that the presence of UL13, but not its kinase activity, is required for efficient accumulation of UL55 in HSV-2-infected cells, whereas UL13 kinase activity is required for efficient accumulation of Us10. Similarly, the CHPK of Marek’s disease herpesvirus has been reported to stabilize Us10 in infected cells (31).

**Fig. 6.**
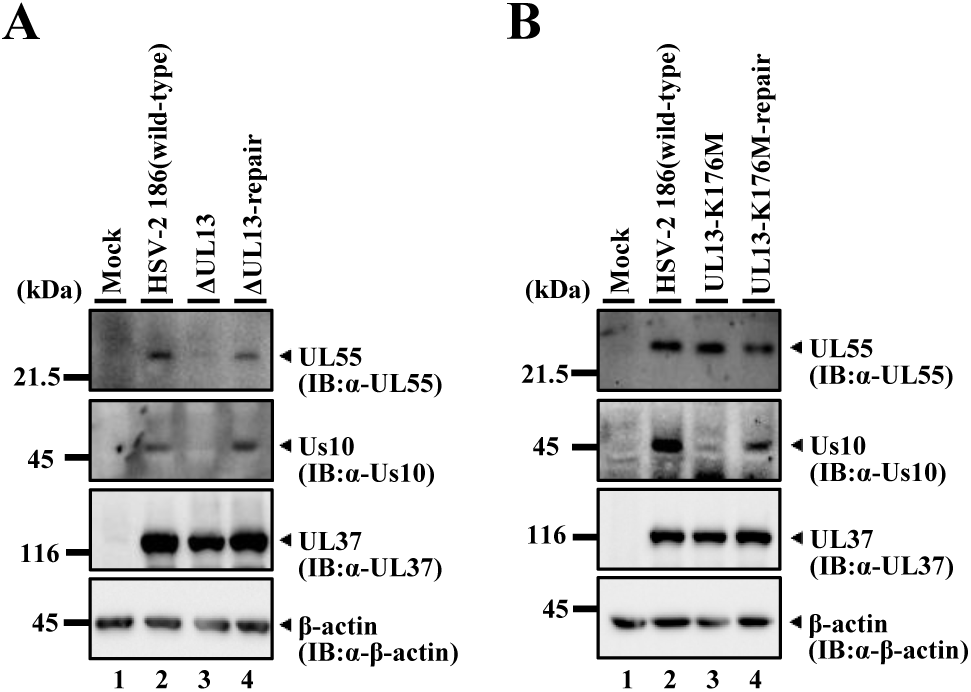
Effects of mutation(s) in UL13 on expression of UL55 and Us10. **A**. Vero cells were mock-infected (lane 1) or infected with wild-type HSV-2 186 (lane 2), YK862 (ΔUL13) (lane 3) or YK863 (ΔUL13-repair) (lane 4) at an MOI of 3, harvested at 24 h post-infection, and lysates were analyzed by immunoblotting with the indicated antibodies. **B.** Vero cells were mock-infected (lane 1) or infected with wild-type HSV-2 186 (lane 2), YK864 (UL13-K176M) (lane 3) or YK865 (UL13-K176M-repair) (lane 4) at an MOI of 3, harvested at 24 h post-infection, and lysates were analyzed by immunoblotting with the indicated antibodies. Digital images are representative of three independent experiments.

### Effects of UL55 and/or Us10 on UL13-mediated phosphorylation of its substrates in HSV-2 infected cells

To examine the effects of UL55 and/or Us10 on UL13 substrates in HSV-2-infected cells, Vero or human osteosarcoma U2OS cells were mock-infected or infected with wild-type HSV-2 186, YK874 (ΔUL55), YK875 (ΔUL55-repair), YK876 (ΔUs10), YK877 (ΔUs10-repair), YK878 (ΔUL55/ΔUs10), YK879 (ΔUL55/ΔUs10-repair) or YK864 (UL13-K176M), lysed and subjected to immunoblotting.

Previous studies have reported that infection with wild-type HSV-2 enhances phosphorylation of EF-1δ at Ser-133, detected by an anti-EF-1δ-S133^P^ monoclonal antibody and leads to the accumulation of the hyperphosphorylated form of EF-1δ, detected as a slower migrating band by immunoblotting with anti-EF-1δ polyclonal antibodies, compared to mock-infection (18). In contrast, such changes are not observed upon infection with YK865 (UL13-K176M) (18). The increase in the hyperphosphorylated form of EF-1δ results from EF-1δ phosphorylation at Ser-133 (4), indicating that enhanced phosphorylation of EF-1δ at this site in HSV-2-infected cells is dependent on UL13 (18). As shown in Fig. 7A, E, and I, EF-1δ phosphorylation at Ser-133 and accumulation of its hyperphosphorylated form in Vero cells infected with YK874 (ΔUL55) were significantly reduced compared to those in cells infected with wild-type HSV-2 186 or YK875 (ΔUL55-repair). In contrast, phosphorylation levels and hyperphosphorylated EF-1δ accumulation in cells infected with YK876 (ΔUs10) were comparable to those in cells infected with wild-type HSV-2 186 or YK877 (ΔUs10-repair) (Fig. 7B, F, and J). Vero cells infected with YK878 (ΔUL55/ΔUs10) exhibited a phosphorylation profile similar to that of YK874 (ΔUL55)-infected cells, although phosphorylation appeared to be further reduced (Fig. 7A, C, E, G, I, and K). Indeed, EF-1δ phosphorylation at Ser-133 and accumulation of its hyperphosphorylated form in cells infected with YK878 (ΔUL55/ΔUs10) were significantly lower than those in cells infected with YK874 (ΔUL55) and were comparable to those observed in cells infected with YK864 (UL13-K176M) (Fig. 8D, H, and L). Similar results were also obtained in U2OS cells (Fig. 8); however, while the difference in the accumulation of hyperphosphorylated EF-1δ between YK878 (ΔUL55/ΔUs10)-and YK864 (UL13-K176M)-infected cells was similarly small as in Vero cells, it was statistically significant only in U2OS cells (Fig. 8L). These results indicate that both UL55 and Us10 are required for optimal UL13 activity to induce phosphorylation of EF-1d at Ser-133 in HSV-2-infected cells.

**Fig. 7.**
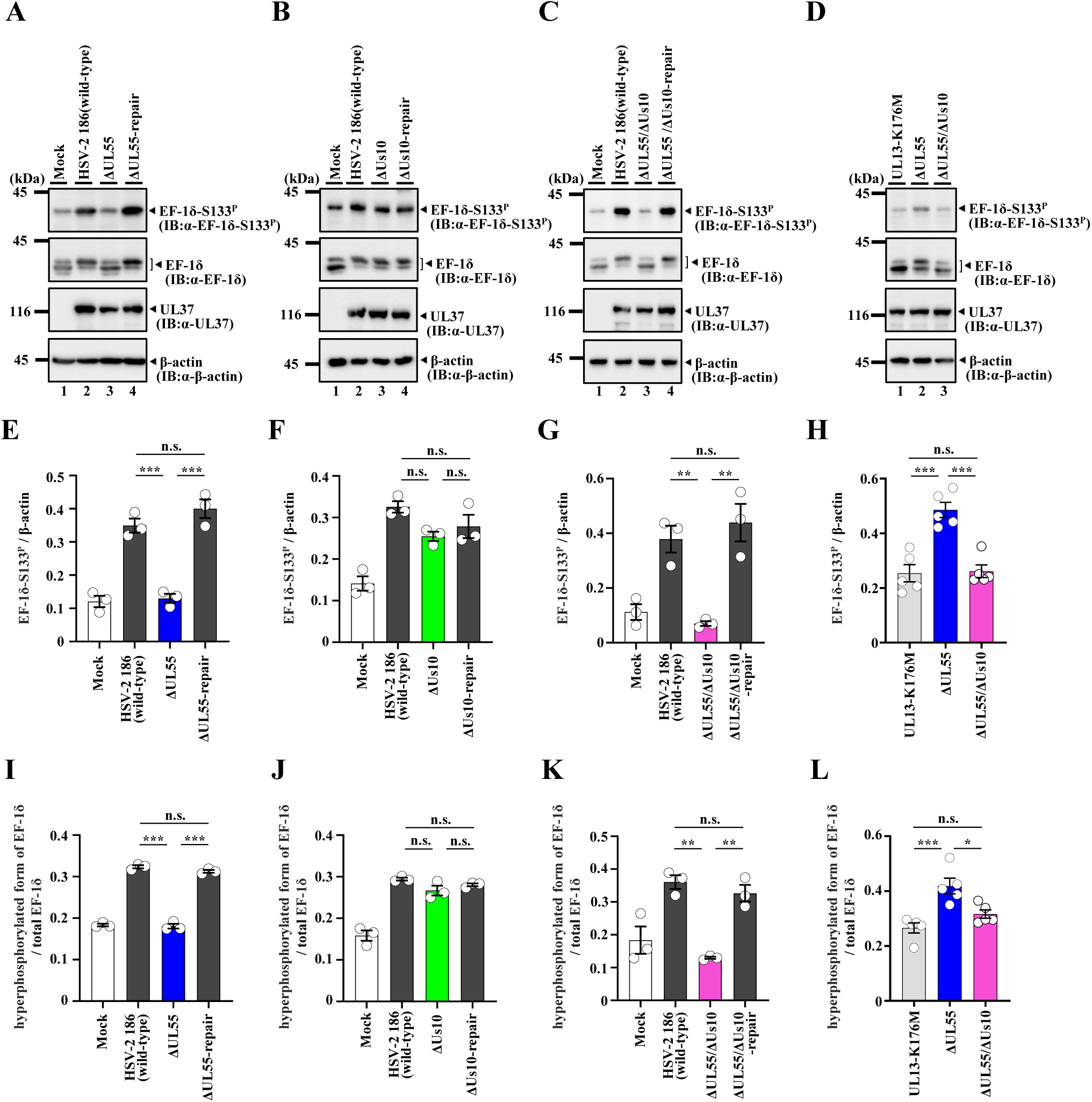
Effects of mutation(s) in UL55 and/or Us10 on phosphorylation of EF-1δ Ser-133 in Vero cells. A-D. U2OS cells were infected with wild-type HSV-2 186 (A-C), YK874 (ΔUL55) (A, D), YK875 (ΔUL55-repair) (A), YK876 (ΔUs10) (B), YK877 (ΔUs10-repair) (B), YK878 (ΔUL55/ΔUs10) (C, D), YK879 (ΔUL55/ΔUs10-repair) (C), or YK864 (UL13-K176M) (D) for 24 h at an MOI of 3 were analyzed by immunoblotting with the indicated antibodies. Digital images are representative of three (A-C) or five (D) independent experiments. **E-H.** Amount of EF-1δ-S133^P^ protein detected with anti-EF-1δ-S133^P^ monoclonal antibody (Fig. 7A-D, top panel) relative to that of β-actin protein detected with anti-β-actin antibody (Fig. 7A-D, bottom panel) in HSV-2-infected cells. Data were normalized by dividing the sum of the data on the same blot (52). Each value is the mean ± SEM of three (E-G) or five (H) experiments. Statistical significance was analyzed by ANOVA with the Tukey’s test. Asterisks indicate statistically significant values (*, *P* < 0.05; **, *P* < 0.01; ***, *P* < 0.001). n.s., not significant. **I-L.** Amount of hyperphosphorylated form of EF-1δ protein detected with anti-EF-1δ polyclonal antibody (Fig. 7A-D, upper band in second panel from the top) relative to that of total EF-1δ protein (hyperphosphorylated and hypophosphorylated forms of EF-1δ) detected with anti-EF-1δ polyclonal antibody (Fig. 7A-D, both bands in second panel from the top) in HSV-2-infected cells. Data were normalized by dividing the sum of the data on the same blot (52). Each value is the mean ± SEM of three (I-K) or five (L) experiments. Statistical significance was analyzed by ANOVA with the Tukey’s test. Asterisks indicate statistically significant values (*, *P* < 0.05; **, *P* < 0.01; ***, *P* < 0.001). n.s., not significant.

**Fig. 8.**
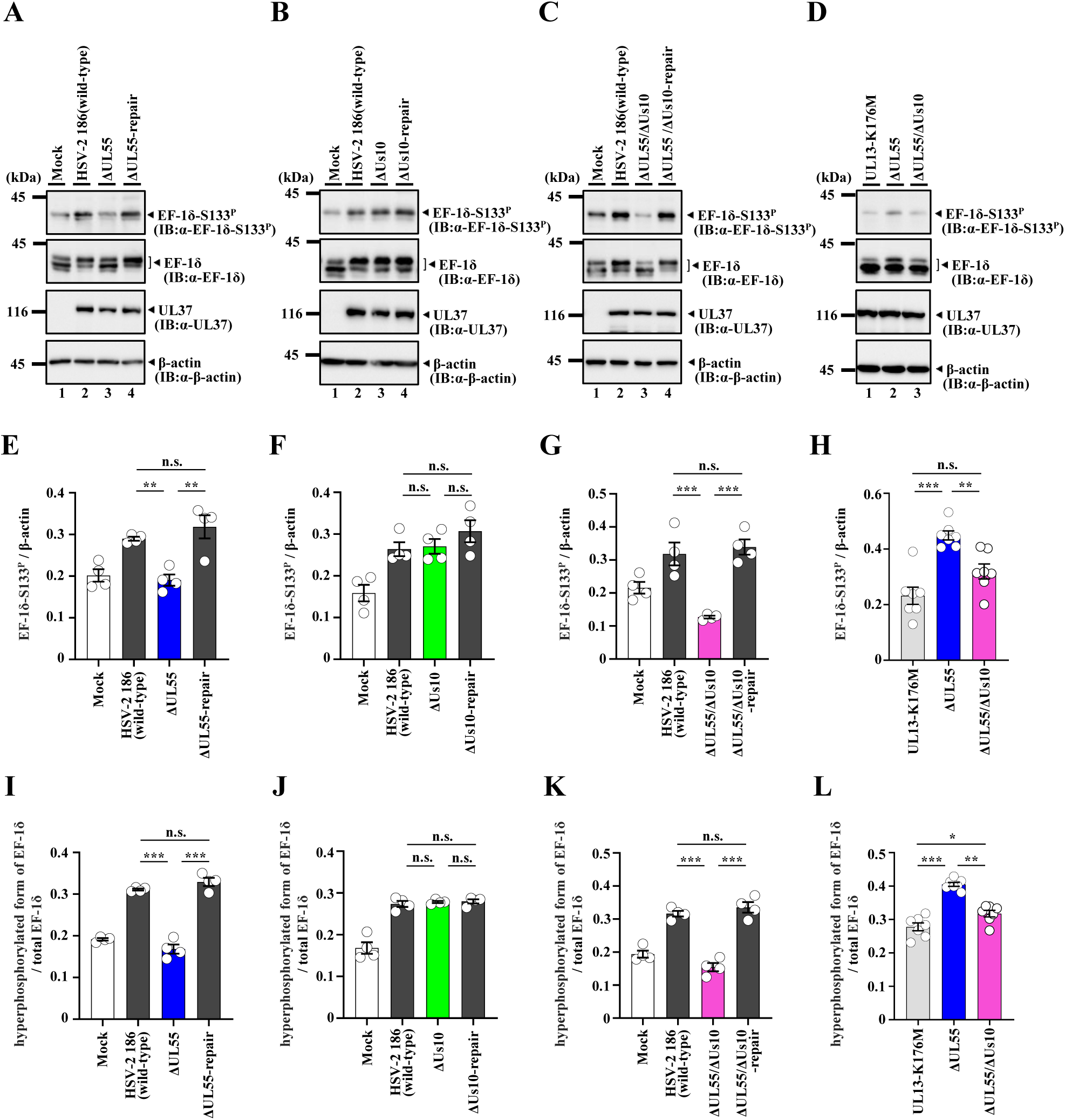
Effects of mutation(s) in UL55 and/or Us10 on phosphorylation of EF-1δ Ser-133 in U2OS cells. A-D. U2OS cells were infected with wild-type HSV-2 186 (A-C), YK874 (ΔUL55) (A, D), YK875 (ΔUL55-repair) (A), YK876 (ΔUs10) (B), YK877 (ΔUs10-repair) (B), YK878 (ΔUL55/ΔUs10) (C, D), YK879 (ΔUL55/ΔUs10-repair) (C), or YK864 (UL13-K176M) (D) for 24 h at an MOI of 3 were analyzed by immunoblotting the indicated antibodies. Digital images are representative of four (A-C) or seven (D) independent experiments. **E-H.** Amount of EF-1δ-S133^P^ protein detected with anti-EF-1δ-S133^P^ monoclonal antibody (Fig. 8A-D, top panel) relative to that of β-actin protein detected with anti-β-actin antibody (Fig. 8A-D, bottom panel) in HSV-2-infected cells. Data were normalized by dividing the sum of the data on the same blot (52). Each value is the mean ± SEM of four (E-G) or seven (H) experiments. Statistical significance was analyzed by ANOVA with the Tukey’s test. Asterisks indicate statistically significant values (*, *P* < 0.05; **, *P* < 0.01; ***, *P* < 0.001). n.s., not significant. **I-L.** Amount of hyperphosphorylated form of EF-1δ protein detected with anti-EF-1δ polyclonal antibody (Fig. 8A-D, upper band in second panel from the top) relative to that of total EF-1δ protein (hyperphosphorylated and hypophosphorylated forms of EF-1δ) detected with anti-EF-1δ polyclonal antibody (Fig. 8A-D, both bands in second panel from the top) in HSV-2-infected cells. Data were normalized by dividing the sum of the data on the same blot (52). Each value is the mean ± SEM of four (I-K) or seven (L) experiments. Statistical significance was analyzed by ANOVA with the Tukey’s test. Asterisks indicate statistically significant values (*, *P* < 0.05; **, *P* < 0.01; ***, *P* < 0.001). n.s., not significant.

**Fig. 9.**
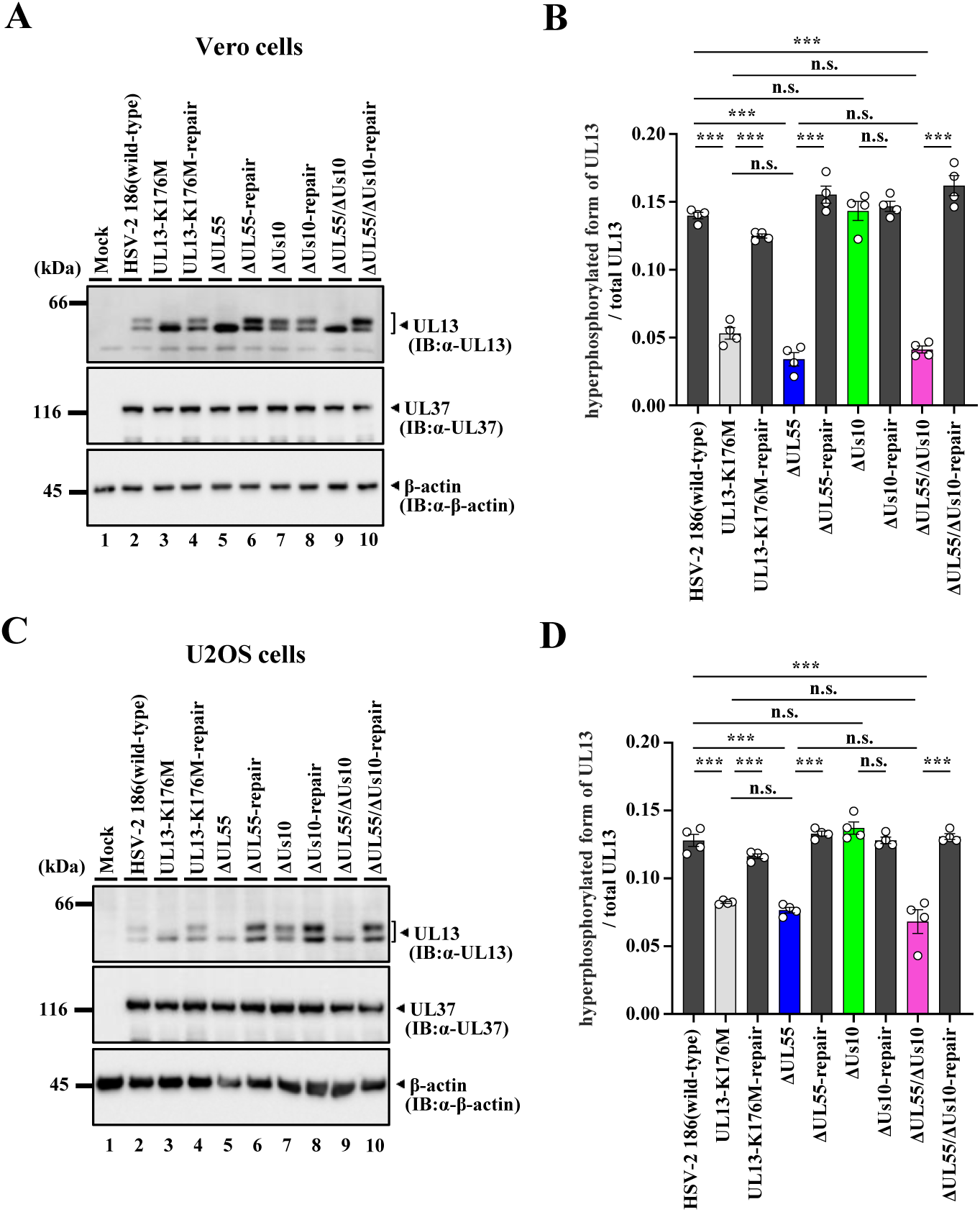
Effects of mutation(s) in UL55 and/or Us10 on expression of UL13 proteins. A, **C.** Vero cells (A) or U2OS cells (C) were mock-infected (lane 1) or infected with wild-type HSV-2 186 (lane 2), YK864 (UL13-K176M) (lane 3), YK865 (UL13-K176M-repair) (lane 4), YK874 (ΔUL55) (lane 5), YK875 (ΔUL55-repair) (lane 6), YK876 (ΔUs10) (lane 7), YK877 (ΔUs10-repair) (lane 8), YK878 (ΔUL55/ΔUs10) (lane 9), or YK879 (ΔUL55/ΔUs10-repair) (lane 10) at an MOI of 3, harvested at 24 h post-infection, and lysates were analyzed by immunoblotting with the indicated antibodies. Digital images are representative of four independent experiments. **B, D.** Amount of hyperphosphorylated form of UL13 protein detected with anti-UL13 monoclonal antibody (Fig. 9A, C, upper band in top panel) relative to that of total UL13 protein (hyperphosphorylated and hypophosphorylated forms of UL13) detected with anti-UL13 monoclonal antibody (Fig. 9A, C, both bands in top panel) in HSV-2-infected cells. Each value is the mean ± SEM of four experiments. Statistical significance was analyzed by ANOVA with the Tukey’s test. Asterisks indicate statistically significant values (***, *P* < 0.0001). n.s., not significant.

Previous studies have reported that auto-phosphorylated UL13 is detected as a slower-migrating band in immunoblotting with an anti-UL13 monoclonal antibody (18). As previously shown (18), UL13 in Vero cells infected with wild-type HSV-2 186 or each of the repaired viruses was detected as two bands with different mobilities (Fig. 9A). In contrast, the slower migrating band corresponding to auto-phosphorylated UL13 was barely detectable in cells infected with YK864 (UL13-K176M) (18). Similarly, auto-phosphorylated UL13 was barely detectable in cells infected with YK874 (ΔUL55) or YK878 (ΔUL55/ΔUs10) (Fig. 9A). The ratio of auto-phosphorylated UL13 to total UL13 in cells infected with YK874 (ΔUL55) was comparable to that in cells infected with YK878 (ΔUL55/ΔUs10) and YK864 (UL13-K176M) (Fig. 9B). In contrast, cells infected with YK876 (ΔUs10) exhibited a phosphorylation profile similar to that of wild-type HSV-2 186-infected cells, and the ratio of auto-phosphorylated UL13 to total UL13 was comparable to that in wild-type HSV-2 186-infected cells (Fig. 9A and B). Similar results were also obtained in U2OS cells (Fig. 9C and D). These results indicate that UL55, but not Us10, is required for optimal UL13 auto-phosphorylation activity in HSV-2-infected cells and are in agreement with our findings above that auto-phosphorylated UL13 was detected only in the presence of UL55 but not Us10 in the *in vitro* kinase assays (Fig. 2B).

### Effects of UL55 and/or Us10 on HSV-2 replication and cell-cell spread

It has been reported that the kinase activity of UL13 is required for efficient HSV-2 replication and cell-cell spread in a manner dependent on multiplicity of infection (MOI) and/or cell type (16, 18). To examine the effects of UL55 and/or Us10 on HSV-2 replication and cell-cell spread in cell cultures, we analyzed progeny virus yields and plaque sizes in U2OS and Vero cells infected with wild-type HSV-2 186, YK864 (UL13-K176M), YK865 (UL13-K176M-repair), YK874 (ΔUL55), YK875 (ΔUL55-repair), YK876 (ΔUs10), YK877 (ΔUs10-repair), YK878 (ΔUL55/ΔUs10), or YK879 (ΔUL55/ΔUs10-repair). As reported previously (18), progeny virus yields of YK864 (UL13-K176M) were significantly lower than those of wild-type HSV-2 186 or YK865 (UL13-K176M-repair) in U2OS cells at an MOI of 0.01, but not at an MOI of 3 (Fig. 10A and B). No significant differences in virus yield were observed in Vero cells at either MOI (Fig. 10C and D). In addition, YK864 (UL13-K176M) formed significantly smaller plaques than wild-type HSV-2 186 or YK865 (UL13-K176M-repair) in U2OS cells, but not in Vero cells (Fig. 10E and F). Similarly, YK874 (ΔUL55) exhibited significantly reduced progeny virus yields in U2OS cells at an MOI of 0.01, comparable to those of YK864 (UL13-K176M), and significantly lower than those of wild-type HSV-2 186 and YK875 (ΔUL55-repair) (Fig. 10A). However, this reduction was not observed at an MOI of 3 in U2OS cells, nor at either MOI in Vero cells (Fig. 10B to D). YK874 (ΔUL55) also produced significantly smaller plaques in U2OS cells, again comparable to those formed by YK864 (UL13-K176M), than those formed by wild-type HSV-2 186 or YK875 (ΔUL55-repair), whereas plaque sizes of these strains were comparable in Vero cells (Fig. 10E and F). In contrast, YK876 (ΔUs10) showed progeny virus yields and plaque sizes comparable to those of wild-type HSV-2 186 and YK877 (ΔUs10-repair) in both U2OS and Vero cells (Fig. 10). The progeny virus yields and plaque sizes of YK878 (ΔUL55/ΔUs10) were similar to those of YK874 (ΔUL55), indicating that deletion of Us10 in addition to UL55 did not further impair viral replication and cell-cell spread under the conditions tested. These results indicate that UL55, but not Us10, is required for efficient HSV-2 replication and cell-cell spread in cell culture, to an extent comparable to that of UL13 kinase activity.

**Fig. 10.**
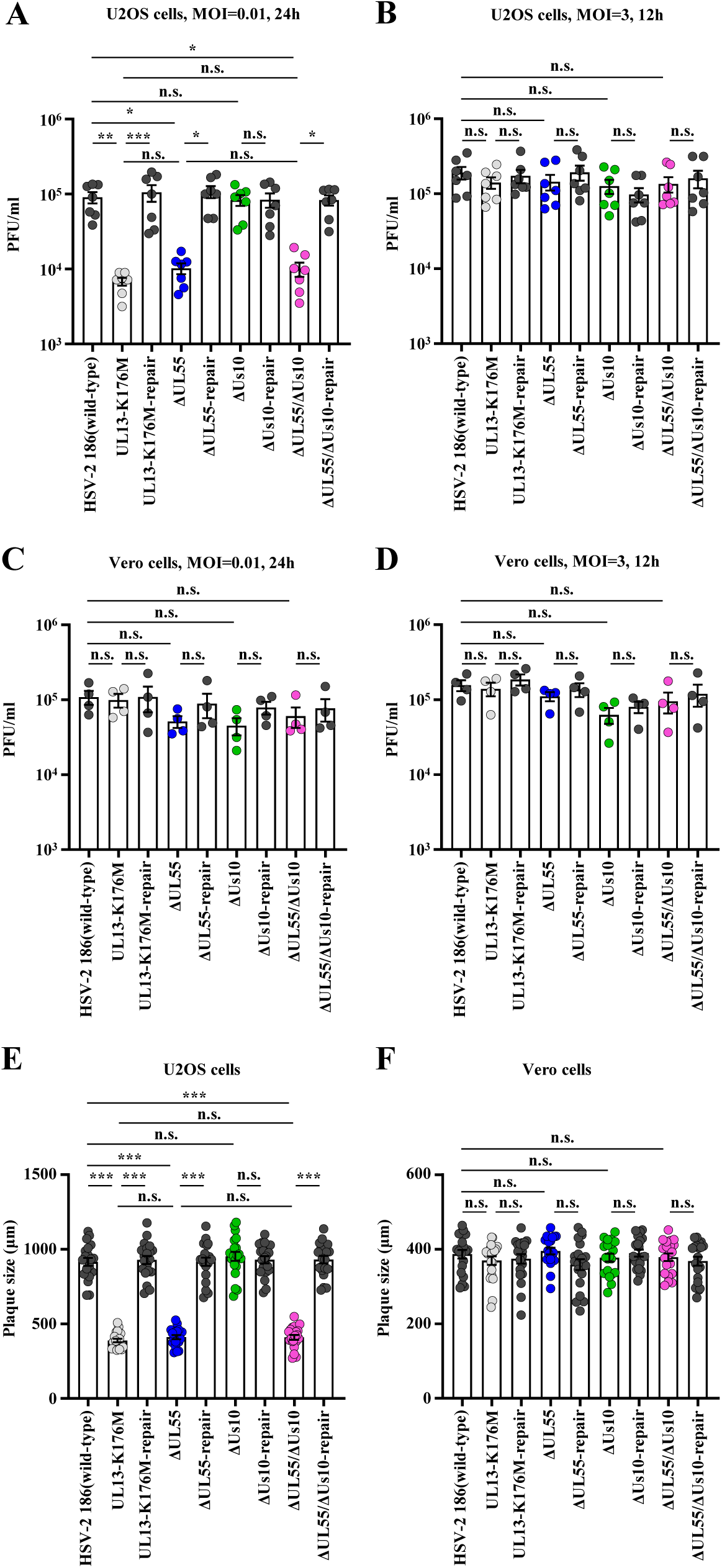
Effects of mutation(s) in UL55 and/or Us10 on viral replication and cell-cell spread in U2OS or Vero cells. A-D. U2OS cells (A, B) or Vero cells (C, D) were infected with wild-type HSV-2 186, YK864 (UL13-K176M), YK865 (UL13-K176M-repair), YK874 (ΔUL55), YK875 (ΔUL55-repair), YK876 (ΔUs10), YK877 (ΔUs10-repair), YK878 (ΔUL55/ΔUs10), or YK879 (ΔUL55/ΔUs10-repair) at an MOI of 0.01 (A, C) or 3 (B, D). Total virus titers in cell culture supernatants and infected cells were harvested at 24 h (A, C) or 12 h (B, D) post-infection and assayed. Each value is the mean ± standard error of the mean (SEM) of seven (A, B) or four (C, D) experiments. Statistical significance was analyzed by ANOVA with the Tukey’s test. Asterisks indicate statistically significant values (*, *P* < 0.05; **, *P* < 0.01; ***, *P* < 0.001). n.s., not significant. **E, F.** U2OS cells (E) or Vero cells (F) were infected with wild-type HSV-2 186, YK864 (UL13-K176M), YK865 (UL13-K176M-repair), YK874 (ΔUL55), YK875 (ΔUL55-repair), YK876 (ΔUs10), YK877 (ΔUs10-repair), YK878 (ΔUL55/ΔUs10), or YK879 (ΔUL55/ΔUs10-repair) at an MOI of 0.0001 under plaque assay conditions. Diameters of 20 single plaques for each virus were measured at 48 h post-infection. Each data point is the mean ± SEM of the measured plaque sizes. Statistical significance was analyzed by ANOVA with Tukey’s test. Asterisks indicate statistically significant values (***, *P* < 0.0001). n.s., not significant. Data are representative of three independent experiments.

### Effects of VZV homologs of HSV-2 UL55 and Us10 on EF-1δ phosphorylation mediated by VZV ORF47

HSV-2 UL55 and Us10 are conserved in the subfamily *Alphaherpesvirinae* (Fig. 11), but not in the *Beta-* and *Gammaherpesvirinae* subfamilies. To investigate whether the effects of HSV-2 UL55 and Us10 on UL13 are conserved in other alphaherpesviruses, we examined whether the VZV homologs of UL55 and Us10, ORF3 and ORF64, respectively, exert similar effects on EF-1δ phosphorylation mediated by VZV ORF47, the homolog of HSV-2 UL13. COS-7 cells were transfected with a plasmid expressing EGFP-EF-1δ(F), together with a plasmid expressing SE-ORF47 or SE-ORF47-K157M, a kinase-dead mutant of ORF47 (3), in combination with a plasmid expressing HA-tagged ORF3 (HA-ORF3) or HA-tagged ORF64 (HA-ORF64), and then subjected to immunoblotting with an anti-EF-1δ-S133^P^ antibody. Phosphorylation of EGFP-EF-1δ(F) at Ser-133 was significantly enhanced by co-expression of SE-ORF47 with HA-ORF3, but not by co-expression of SE-ORF47-K157M with HA-ORF3 (Fig. 12A and B). In contrast, co-expression of SE-ORF47 with HA-ORF64 had little effect on phosphorylation of EGFP-EF-1δ(F) at Ser-133 (Fig. 12C, D). These results indicate that VZV ORF3, but not ORF64, can enhance the ability of ORF47 to phosphorylate EF-1δ at Ser-133.

**Figure 11.**
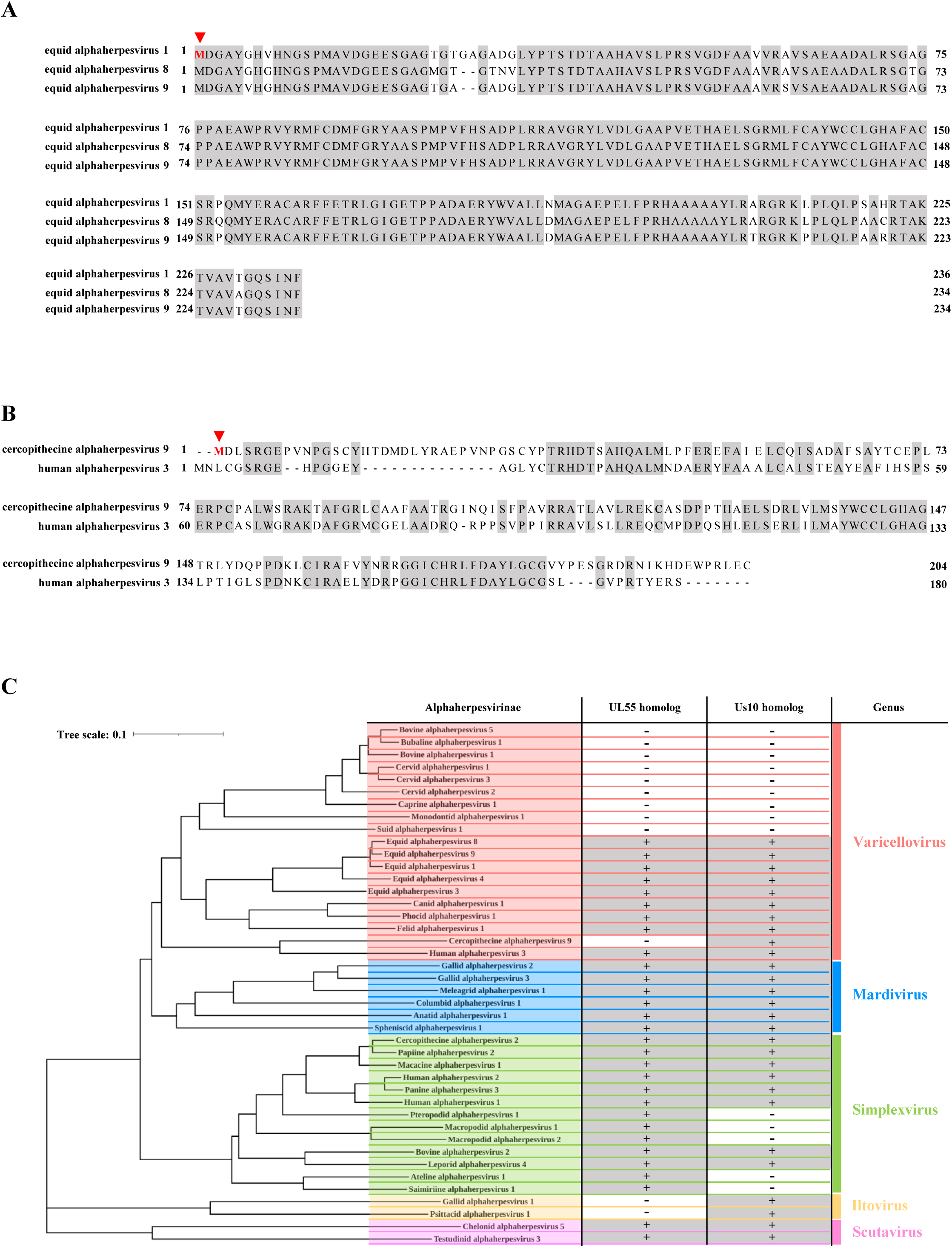
Conservation and phylogenetic distribution of HSV-1 UL55 and Us10 homologs among *alphaherpesviruses*. A, **B.** Multiple-sequence alignments of Us10 homologs from phylogenetically adjacent taxa (see panel C). Multiple-sequence alignment of annotated ORF4 proteins (Us10 homologs) from *Equid alphaherpesvirus 8* and *Equid alphaherpesvirus 9* together with the as-yet-unannotated Us10 homolog encoded by *Equid alphaherpesvirus 1* (A). Multiple-sequence alignment of annotated ORF3 proteins (Us10 homologs) from *Human alphaherpesvirus 3* aligned with the as-yet-unannotated Us10 homolog of *Cercopithecine alphaherpesvirus 9* (B). Residues conserved in every sequence are shaded black; methionines that may serve as translational start codons are colored red and indicated with red arrowheads. These alignments indicate the presence of Us10 homolog genes in Equid alphaherpesvirus 1and Cercopithecine alphaherpesvirus 9. **C.** Phylogenetic tree of *alphaherpesviruses* inferred from concatenated amino-acid sequences of six core genes conserved throughout the family *Herpesviridae*—uracil-DNA glycosylase, helicase-primase helicase subunit, DNA-packaging terminase subunit 1, major capsid protein, envelope glycoprotein B, and DNA polymerase catalytic subunit. The topology was visualized in iTOL (https://itol.embl.de/) after importing a pre-computed Newick tree (https://ictv.global/sites/default/files/report_files/Alpha_Feb21_treefile.txt) obtained from the ICTV Online Report, Subfamily *Alphaherpesvirinae* (https://ictv.global/report/chapter/orthoherpesviridae/orthoherpesviridae/alphaherpesvirinae); licensed under CC BY-SA 4.0). A table to the right of each taxon denotes the presence (+) or absence (–) of homologs of HSV-1 UL55 and Us10, as determined from NCBI annotations or alignment-based curation in panels (A) and (B). Accession numbers and corresponding NCBI hyperlinks are listed in Table S1.

**Fig. 12.**
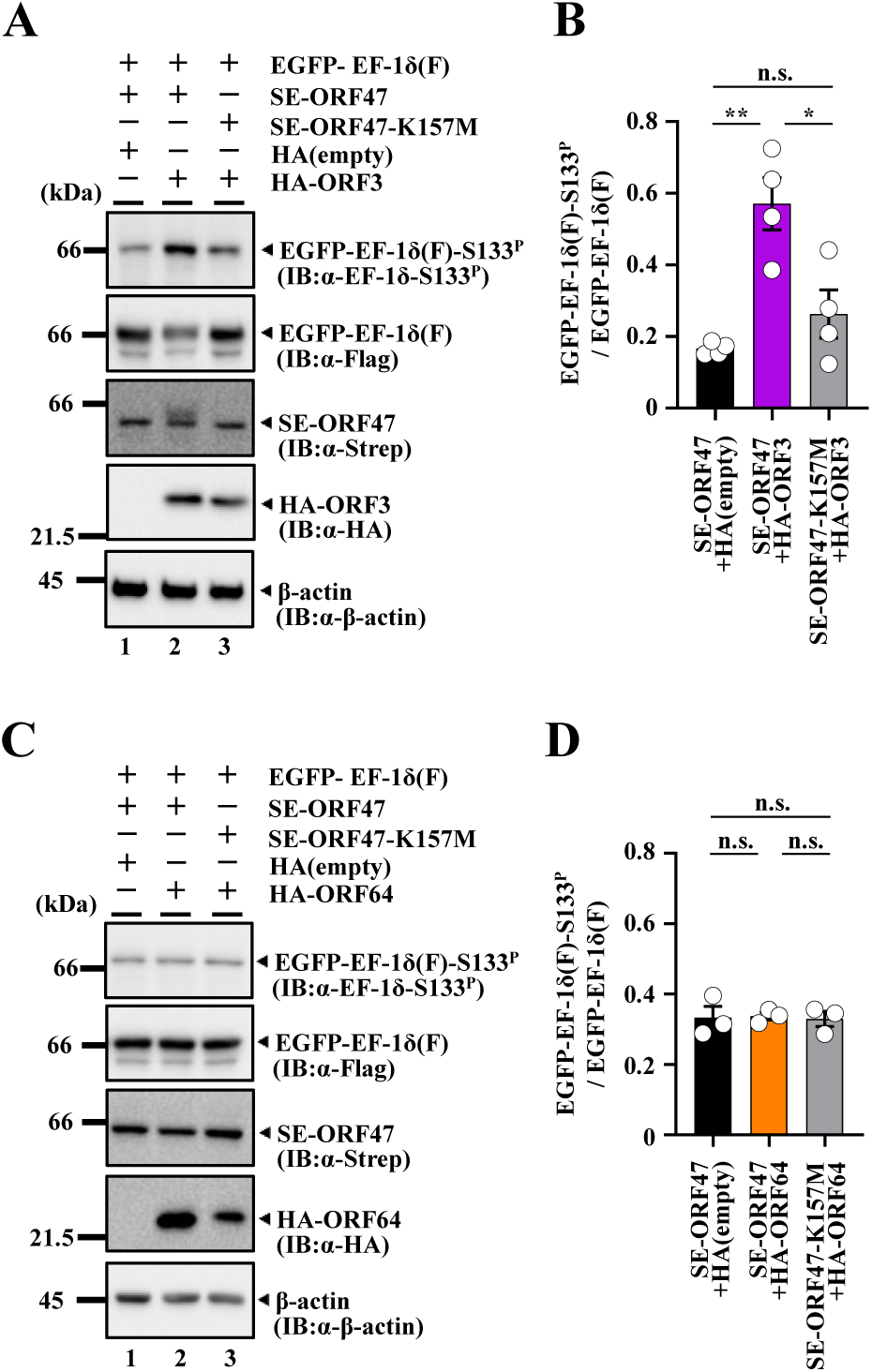
VZV ORF3 interacts with ORF47 and functions as an activator of ORF47. **A.** COS-7 cells were transfected with a plasmid expressing EGFP-EF-1δ(F) in combination with a plasmid expressing SE-ORF47 (lanes 1, 2) or SE-ORF47-K157M (lane 3), and an empty plasmid (lane 1) or a plasmid expressing HA-ORF3 (lanes 2, 3), harvested 48 h post-transfection, and lysates were then analyzed by immunoblotting with the indicated antibodies. Digital images are representative of four independent experiments. **B.** Amount of EGFP-EF-1δ(F)-S133P protein detected with anti-EF-1δ-S133P monoclonal antibody (Fig. 12A, top panel) relative to that of EGFP-EF-1δ(F) protein detected with anti-Flag antibody (Fig. 12A, second panel from the top) in transfected cells. Data were normalized by dividing the sum of the data on the same blot (52). Each value is the mean ± SEM of four experiments. Statistical significance was analyzed by ANOVA with Tukey’s test. Asterisks indicate statistically significant values (*, *P* < 0.05; **, *P* < 0.01). n.s., not significant. **C.** COS-7 cells were transfected with a plasmid expressing EGFP-EF-1δ(F) in combination with a plasmid expressing SE-ORF47 (lanes 1, 2) or SE-ORF47-K157M (lane 3), and an empty plasmid (lane 1) or a plasmid expressing HA-ORF64 (lanes 2, 3), harvested 48 h post-transfection, and lysates were then analyzed by immunoblotting with the indicated antibodies. Digital images are representative of three independent experiments. **D.** Amount of EGFP-EF-1δ(F)-S133P protein detected with anti-EF-1δ-S133P monoclonal antibody (Fig. 12C, top panel) relative to that of EGFP-EF-1δ(F) protein detected with anti-Flag antibody (Fig. 12C, second panel from the top) in transfected cells. Data were normalized by dividing the sum of the data on the same blot (52). Each value is the mean ± SEM of three experiments. Statistical significance was analyzed by ANOVA with Tukey’s test. n.s., not significant.

## Discussion

In this study, we identified HSV-2 UL55 and Us10 as viral activators of UL13. Co-expression of UL13 with either UL55 or Us10 significantly enhanced phosphorylation of EF-1δ at Ser-133 compared to the expression of UL13 alone. Moreover, UL13 was co-precipitated with either UL55 or Us10 upon co-expression, and its kinase activity was substantially increased in the presence of either of these viral proteins. These findings suggest that UL55 and Us10 interact with UL13 and act as activators of this viral protein kinase. In HSV-2-infected cells, UL13 was specifically co-precipitated with UL55 and Us10. Notably, the UL55-null mutation markedly reduced phosphorylation of EF-1δ at Ser-133, whereas the Us10-null mutation had little effect. However, the double-null mutation further decreased the phosphorylation compared to the UL55-null mutation alone. Similarly, UL13 autophosphorylation was reduced by the UL55-null mutation, while the Us10-null mutation had minimal effect. However, no further reduction was observed in the double-null mutant compared to the UL55-null mutant. In agreement with this, auto-phosphorylated UL13 was detected only in the presence of UL55, but not Us10, *in vitro*. These results suggest that UL55 acts as a principal activator of UL13 in HSV-2-infected cells, while Us10 serves as an auxiliary activator. To our knowledge, the functions of HSV UL55 and Us10 in infected cells have remained unclear, and this study represents the first to elucidate their functional roles.

We showed that the UL55-null mutation reduced UL13 autophosphorylation to a level comparable to that observed with the kinase-dead mutation in UL13 in HSV-2-infected cells, and that individually expressed UL13 in COS-7 cells was unable to induce phosphorylation of EF-1δ at Ser-133. In contrast, previous studies have reported that highly purified UL13, when expressed individually using a baculovirus expression system, has intrinsic kinase activity capable of both auto-phosphorylation and substrate trans-phosphorylation *in vitro* (4). Collectively, these observations suggest that, while UL55 is not essential for the intrinsic kinase activity of UL13, it is crucial for the functional expression of UL13 activity in HSV-2-infected cells. In agreement with this, the UL55-null mutation reduced HSV-2 replication and cell-cell spread in U2OS cells to levels comparable to those observed with the kinase-dead mutation in UL13, suggesting that the role of the UL13 kinase activity in HSV-2 replication and cell-cell spread in these cells is largely dependent on UL55.

We demonstrated that individual expression of VZV CHPK ORF47 was insufficient to induce phosphorylation of EF-1δ at Ser-133, whereas co-expression of VZV ORF47 with ORF3, the HSV-2 UL55 homolog, but not with ORF64, the HSV-2 Us10 homolog, significantly enhanced phosphorylation of EF-1δ at this site. These findings suggest that the function of HSV-2 UL55 in activating UL13 is conserved in VZV ORF3, and potentially in other alphaherpesvirus UL55 homologs. In agreement with this, it has been reported that VZV ORF47 mediates phosphorylation of IRF3 in VZV-infected cells, whereas individual expression of ORF47 alone fails to do so, suggesting that a viral cofactor(s) is required for ORF47-mediated phosphorylation of IRF3 in VZV-infected cells (32). Although our results do not support a role for VZV ORF64 as a coactivator of ORF47, we cannot entirely rule out this possibility, given that HSV-2 Us10 exhibited only modest coactivator activity. Further studies will be necessary to clarify whether ORF64 contributes to the regulation of ORF47. As shown in Fig. 11, alphaherpesviruses can be categorized into three groups based on conservation of UL55 and/or Us10 homologs: (i) viruses encoding both UL55 and Us10 such as HSV-1 (human alphaherpesvirus 1) and HSV-2 (human alphaherpesvirus 2); (ii) viruses lacking both genes such as suid alphaherpesvirus 1 (pseudorabies virus, PRV) and bovine alphaherpesvirus 1; and (iii) viruses encoding only one of the two such as saimiriine alphaherpesvirus 1 and gallid alphaherpesvirus 1. In contrast to the CHPKs of HSV-2 and VZV that require viral cofactors for activation, PRV UL13 appears to possess sufficient intrinsic kinase activity without viral cofactors, effectively phosphorylating its substrates when expressed alone in mammalian cells, similarly to what is observed in PRV-infected cells (33, 34), despite the absence of UL55 and Us10 homologs. These observations suggest that PRV UL13 has evolved self-sufficient activation mechanisms, comparable to those of CHPKs in beta-and gammaherpesviruses. Notably, phylogenic analyses based on amino acid sequences of six genes of alphaherpesviruses conserved thoughout the family *Herpesviridae* revealed that viruses lacking both UL55 and Us10 genes form a monophyletic clade within the *Varicellovirus* genus, while viruses encoding Us10 but not UL55 cluster in the *Itovirus* genus (Fig. 11). By contrast, viruses encoding UL55 but not Us10 form two distinct monophyletic clades within the *Simplexvirus* genus (Fig. 11). These phylogenetic patterns suggest that the gain or loss of specific CHPK cofactors might have occurred independently in different alphaherpesvirus lineages, potentially as part of their adaptation to distinct host environments and replication strategies. Taken together, these observations suggest that alphaherpesvirus CHPKs have evolved diverse regulatory strategies, ranging from intrinsic activity to reliance on specific viral cofactors such as UL55 and Us10. This mechanistic diversity is mirrored in their phylogenetic separation, offering new insight into the evolutionary trajectories and functional specialization of CHPK regulation among herpesviruses.

At present, the precise mechanisms by which HSV-2 UL55 and Us10 activate UL13 remain to be elucidated. Protein kinases are known to be regulated by a variety of cofactor proteins that modulate their activity through diverse mechanisms, including conformational changes, promotion of dimerization or oligomerization, subcellular localization, substrate recruitment, or stabilization of the kinase itself (35, 36). Notably, our study revealed a reciprocal regulatory relationship between these viral proteins. Specifically, although the kinase activity of UL13 was not required for UL55 accumulation, the absence of UL13 protein resulted in markedly reduced UL55 accumulation in HSV-2-infected cells, suggesting that UL13 stabilizes UL55 in a kinase-independent manner. In contrast, Us10 accumulation was markedly reduced in the absence of UL13 kinase activity, suggesting that UL13-mediated phosphorylation may contribute to Us10 stabilization in infected cells. These bidirectional regulatory relationships appear to parallel the activation and stabilization dynamics observed in CDK-cyclin systems, in which cyclins activate CDKs and, in turn, are stabilized through binding to CDKs (37–39), with their stability further modulated by CDK phosphorylation (38). UL13 and other CHPKs are classified as CDK-like kinases, as they mimic both the substrate specificity and negative regulatory mechanisms of CDKs (4–9, 12–16). Our data raise the intriguing possibility that HSV-2 UL55 and/or Us10 might function in a manner analogous to cyclins. Binding of cyclins to CDKs induces a conformational change in the CDK activation loop, leading to kinase activation (39). Further studies, including structural and biochemical analyses, will be necessary to clarify the mechanisms by which UL55 and Us10 activate UL13. In particular, investigating whether UL55 or Us10 binding induces conformational changes in the activation loop of UL13 would be of great interest.

## Materials and Methods

### Cells and viruses

Simian kidney epithelial Vero and COS-7 cells, rabbit skin cells, human osteosarcoma U2OS cells, human embryonic kidney epithelial HEK293T cells, and HSV-2 wild-type strain HSV-2 186 were described previously (4, 40–43). Recombinant virus HSV-2 ΔUL13 (YK862) in which the UL13 gene was disrupted by deleting UL13 codons 159-417, recombinant virus HSV-2 ΔUL13-repair (YK863) in which the UL13-null mutation was repaired, recombinant virus HSV-2 UL13-K176M (YK864) encoding an enzymatically inactive UL13 mutant in which lysine at UL13 residue 176 was replaced with methionine, recombinant virus HSV-2 UL13-K176M-repair (YK865) in which the K176M mutation was repaired, were described previously (18) (Fig. 3).

### Plasmids

pcDNA-SE-UL13 or pcDNA-SE-UL13-K176M were constructed by amplifying the entire coding sequence of HSV-2 UL13 from pYEbac861 or the UL13-K176M genome, respectively, by PCR using the primers listed in Table 1, and cloning it into pcDNA-SE (16, 44) in frame with a Strep-tag sequence. pcDNA-SE-ORF47, pcDNA-SE-UL97, pcDNA-SE-BGLF4, and pcDNA-SE-ORF36 were constructed by amplifying the entire coding sequence of VZV ORF47 from pOka BAC DNA (a generous gift from Y. Mori), HCMV UL97 from cDNA synthesized from the total RNA of HEL cells infected with HCMV ADsubUL21.5 (a generous gift from T. Shenk), EBV BGLF4 from pME-BGLF4 (20), or KSHV ORF36 from KSHV DNA isolated from BJAB-BAC36 cells (a generous gift from K. Ueda), respectively, by PCR using the primers listed in Table 1, and cloning it into pcDNA-SE (16) in frame with a Strep-tag sequence. pcDNA-SE-ORF47-K157M, in which Lys-157 of ORF47 was replaced with methionine, was constructed by PCR from pcDNA-SE-ORF47 using the primers listed in Table 1, and cloning it into pcDNA-SE (16) in frame with a Strep-tag sequence as described previously (45). pcDNA-SE-U69 was described previously (16).

**Table. 1.**
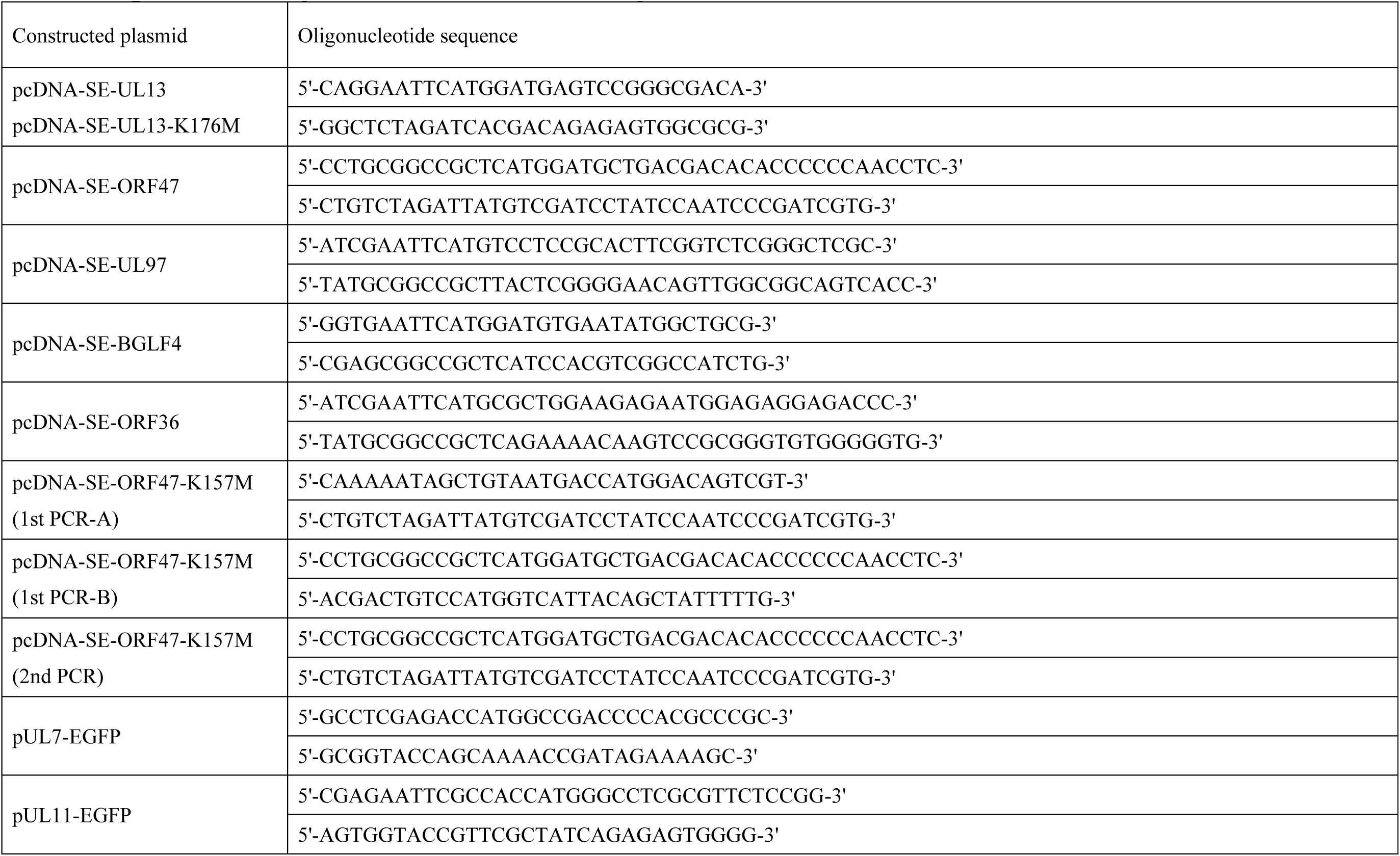
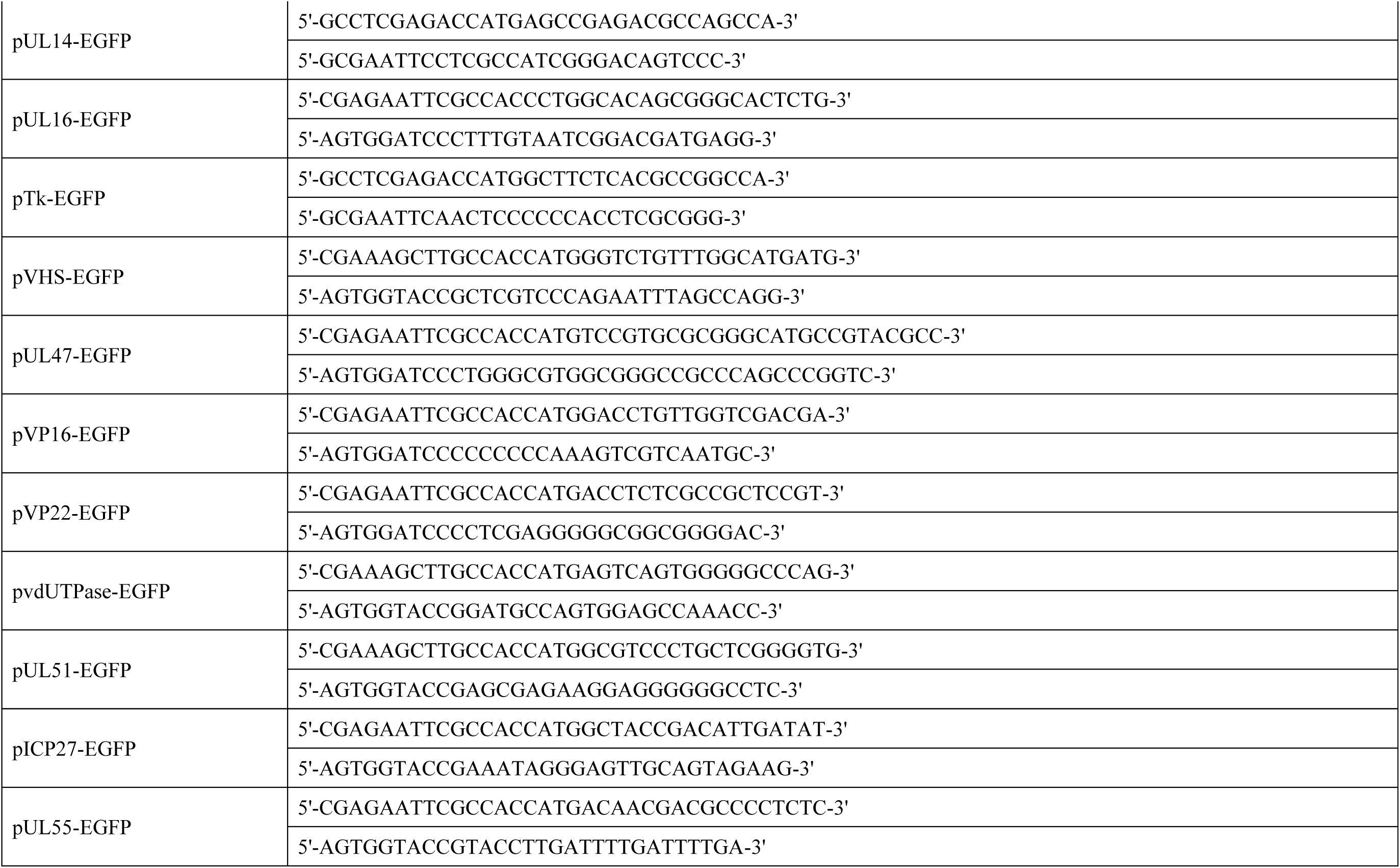
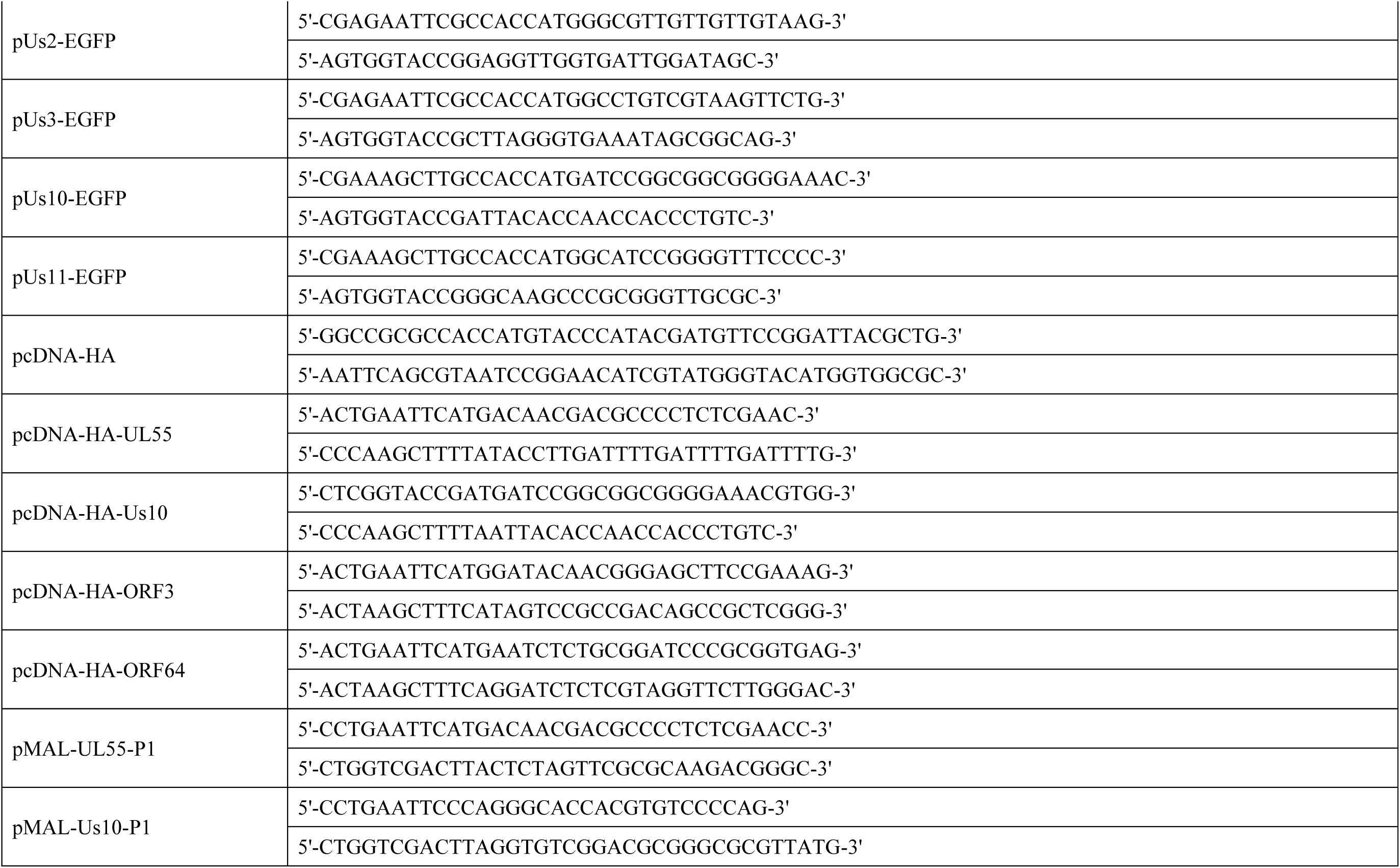
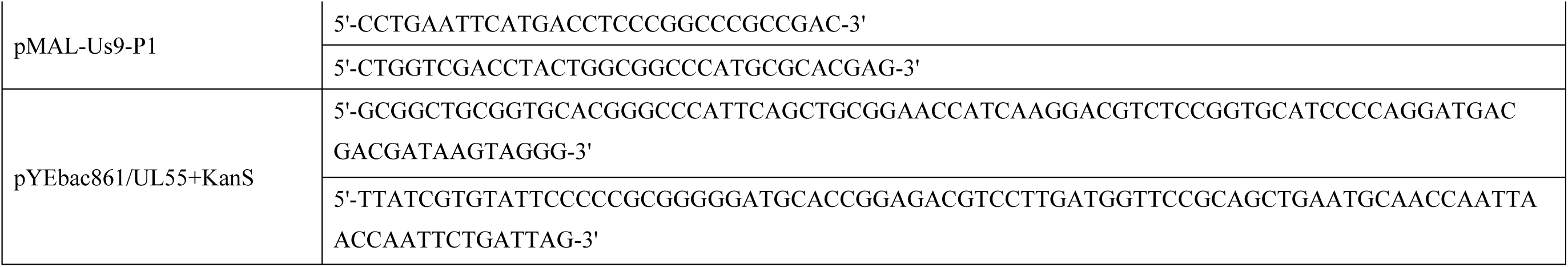
Oligonucleotide sequences for the construction of plasmids.

pUL7-EGFP, pUL11-EGFP, pUL14-EGFP, pUL16-EGFP, pTk-EGFP, pVHS-EGFP, pUL47-EGFP, pVP16-EGFP, pVP22-EGFP, pvdUTPase-EGFP, pUL51-EGFP, pICP27-EGFP, pUL55-EGFP, pUs2-EGFP, pUs3-EGFP, pUs10-EGFP, or pUs11-EGFP were constructed by amplifying the entire coding sequence of each HSV-2 ORF from pYEbac861 by PCR using the primers listed in Table 1, and cloning it into pEGFP-N2 (Clontech) in frame with the EGFP sequence. Based on the genome information of HSV-2 strain HG52, pUL37-EGFP, pICP0-EGFP, pICP4-EGFP, or pICP34.5-EGFP were synthesized by GenScript. pUL36-EGFP was constructed based on the genomic sequence of HSV-2 strain HG52 by synthesizing the N-terminal region of UL36 and cloning it in-frame with the EGFP sequence in the pEGFP-N2 vector (Clontech) by GenScript, followed by insertion of the synthesized C-terminal region of UL36 between the N-terminal region of UL36 and the EGFP sequence. pEGFP-EF-1δ(F), in which EF-1δ was tagged with the Flag epitope and EGFP, as described previously (18). pcDNA-HA, a hemagglutinin (HA) tag with an influenza virus HA epitope, was constructed by annealing the oligonucleotides listed in Table 1 and cloning it into pcDNA3.1/myc-His(-) A (Thermo Fisher Scientific). pcDNA-HA-UL55 or pcDNA-HA-Us10 were constructed by amplifying the entire coding sequence of HSV-2 UL55 or Us10 from pYEbac861 genome by PCR using the primers listed in Table 1, and cloning it into pcDNA-HA in frame with an HA-tag sequence. pcDNA-HA-ORF3 or pcDNA-HA-ORF64 were constructed by amplifying the entire coding sequence of VZV ORF3 or ORF64 from pOka BAC genome by PCR using the primers listed in Table 1, and cloning it into pcDNA-HA in frame with an HA-tag sequence.

pMAL-UL55-P1, pMAL-Us10-P1, or pMAL-Us9-P1 were constructed by amplifying the domains of HSV-2 UL55 (encoded by UL55 codons 1 to 100), Us10 (encoded by Us10 codons 33 to 132), or Us9 (encoded by Us9 codons 1 to 50) from pYEbac861 by PCR using the primers listed in Table 1, and cloning it into pMAL-c (New England BioLabs) in frame with MBP. pMAL-EF-1δ(107–146) or pMAL-EF-1δ(107–146)-S133A were described previously (4).

To construct the transfer plasmid pYEbac861/UL55+KanS, used for generating recombinant viruses YK875 (ΔUL55-repair) and YK879 (ΔUL55/ΔUs10-repair) in which the UL55-null mutation in YK874 (ΔUL55) and the UL55/Us10-null mutations in YK878 (ΔUL55/ΔUs10), respectively, were repaired (Fig. 3), linear fragments containing a gene encoding the I-SceI site, kanamycin resistance, and 83 bp of UL55 sequences were generated by PCR using the primers listed in Table 1 with pEP-KanS (46) as the template. The linear PCR-generated fragments were electroporated into the electrocompetent cells of *Escherichia coli* (*E. coli*) GS1783 containing pYEbac861 (18). These bacteria were then plated on LB agar plates containing 20 μg/ml of chloramphenicol and 40 μg/ml of kanamycin to select *E. coli* clones harboring the kanamycin resistance gene inserted into the UL55 locus. After 36 h, kanamycin-resistant colonies were screened by PCR with appropriate primers, which led to the identification of pYEbac861/UL55+KanS, a *E. coli* GS1783 strain harboring the HSV-2-BAC plasmid pYEbac861/UL55+KanS.

### Mutagenesis of viral genomes and generation of recombinant HSV-2

Recombinant virus YK873 (UL13-HA), carrying a HA-tag at the C-terminus of the UL13 gene in YK861 (UL13-WT) (18) (Fig. 3), was constructed by the two-step Red-mediated mutagenesis procedure using *E. coli* strain GS1783 containing pYEbac861 (18), as described previously (46, 47), except using the primers listed in Table 2. Recombinant virus YK874 (ΔUL55), in which the entire UL55 ORF was deleted (Fig. 3); and YK876 (ΔUs10), in which a tyrosine residue at the position 14 in Us10 was replaced with a stop codon (Fig. 3), were generated by the two-step Red-mediated mutagenesis procedure using *E. coli* strain GS1783 containing pYEbac861 (18), as described previously (46, 47), except using the primers listed in Table 2. Recombinant virus YK878 (ΔUL55/ΔUs10), carrying both the UL55 and Us10 mutations found in YK874 (ΔUL55) and YK876 (ΔUs10) (Fig. 3), was constructed by the two-step Red-mediated mutagenesis procedure using *E. coli* GS1783 carrying the YK874 (ΔUL55) genome as described previously (46, 47), except using the primers listed in Table 2. Recombinant virus YK877 (ΔUs10-repair), in which the mutations in YK876 (ΔUs10) was repaired (Fig. 3), was generated as described previously (46, 47), except using the primers listed in Table 2. For generation of recombinant virus YK875 (ΔUL55-repair), in which the mutation in YK874 (ΔUL55) was repaired (Fig. 3), was generated as described previously (46, 47), except using the primers listed in Table 2, pYEbac861/UL55+KanS, and *E.coli* GS1783 containing the YK874 (ΔUL55) genome (Fig. 3). Recombinant virus YK879 (ΔUL55/ΔUs10-repair), in which the mutations in YK878 (ΔUL55/ΔUs10) were repaired (Fig. 3), was generated by sequentially applying the repair procedures used for YK875 (ΔUL55-repair) and YK877 (ΔUs10-repair).

**Table 2.**
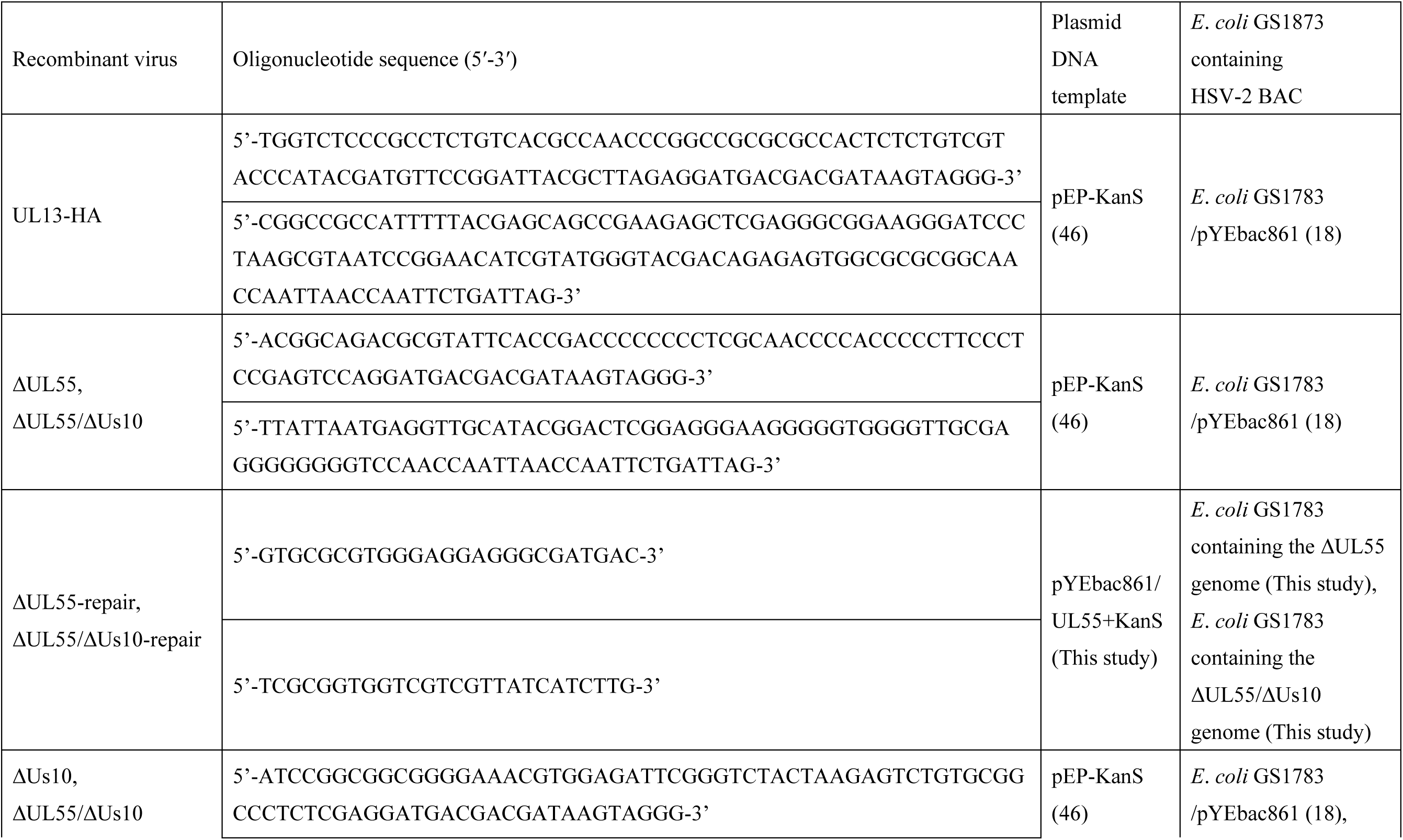
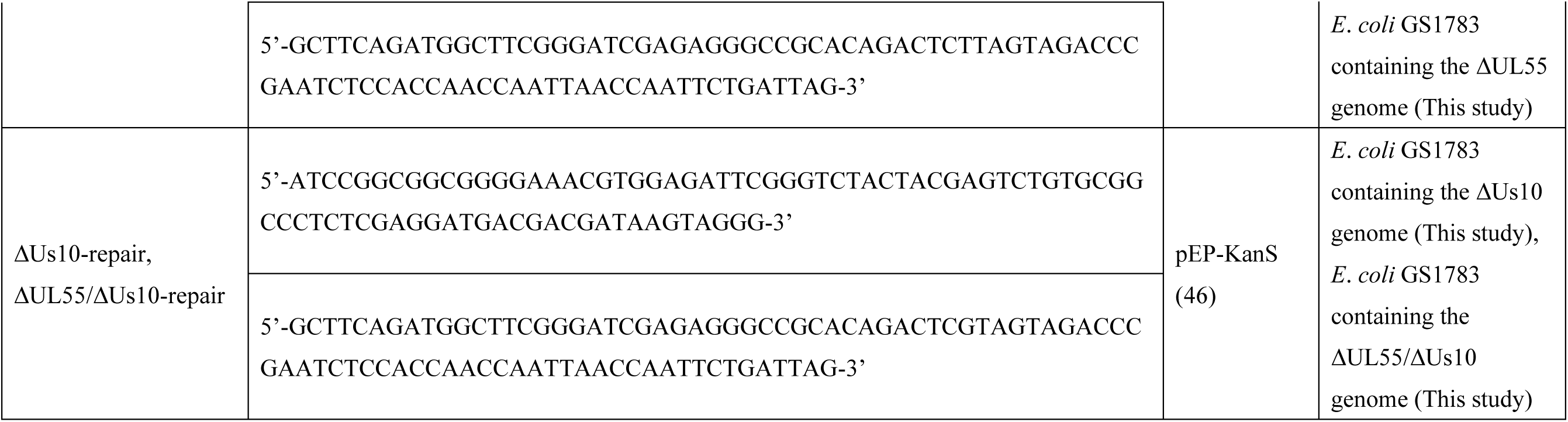
Oligonucleotide sequences for the construction of recombinant viruses.

### Production and purification of MBP fusion proteins

MBP-UL55-P1, MBP-Us10-P1, and MBP-Us9-P1 were expressed in *E. coli* Rosetta (Novagen), transformed with pMAL-UL55-P1, pMAL-Us10-P1, or pMAL-Us9-P1 purified using amylose beads (New England Biolabs), respectively, as described previously (4). The MBP fusion proteins were eluted with MBP elution-buffer (50 mM Tris-HCl [pH 8.0], 25 mM EGTA, 10 mM D(+)-Maltose Monohydrate) and stored at −80 °C. MBP-EF-1δ(107–146) or MBP-EF-1δ(107–146)-S133A were expressed in *E. coli* Rosetta (Novagen), transformed with pMAL-EF-1δ(107–146) or pMAL-EF-1δ(107–146)-S133A purified amylose beads (New England Biolabs), respectively, as described previously (4).

### Antibodies

Antibodies and dilutions used in immunoblotting were as follows: commercial mouse monoclonal antibodies to Flag-tag (M2; Sigma, 1:1000), Strep-tag II (4F1; MBL, 1:1000), HA-tag (TANA2; MBL, 1:1000), β-actin (AC15; Sigma, 1:1000), ICP27 (H1142; Santa Cruz, 1:2000) and rabbit polyclonal antibodies to VP23 (CAC-CT-HSV-UL18; Cosmo Bio, 1:2000), UL37 (CAC-CT-HSV-UL37; CosmoBio, 1:2000) green fluorescent protein (GFP) (598; MBL; 1:1000). Mouse monoclonal antibodies to UL13 (1:500) and EF-1δ with phosphorylated Ser-133 (1:5000) and rabbit polyclonal antibodies to UL56 (1:2000), Us11 (1:500), and EF-1δ (1:1000) were reported previously (18, 48–51). To generate mouse polyclonal antibodies to HSV-2 UL55, Us10 or Us9, BALB/c mice were immunized once with purified MBP-UL55-P1, MBP-Us10-P1 or MBP-Us9-P1, respectively, in combination with TiterMax Gold (TiterMax USA, Inc.). Sera from immunized mice were used as sources of mouse polyclonal antibodies to UL55 (1:100), Us10 (1:100), or Us9 (1:100).

### Transfection

COS-7 or 293T cells were transfected with the indicated plasmid(s) using PEI Max (Polysciences).

### Immunoblotting

Immunoblotting was performed as described previously (51). Brightness/contrast of raw blots were equally adjusted across the entire image with Image lab software (BioRad) to generate representative images. The amount of protein in immunoblot bands was quantitated using ChemiDoc MP (Bio-Rad) with Image Lab 6.1.0 software (Bio-Rad) according to the manufacturer’s instructions.

### *In vitro* kinase assays

HEK293T cells were transfected with SE-UL13 or SE-UL13-K176M in combination with pEGFP-N2, UL55-EGFP or Us10-EGFP. Transfected cells were harvested at 48 h post-transfection and lysed in 0.1% NP-40 buffer (50 mM Tris-HCl [pH 8.0], 150 mM NaCl, 50 mM NaF, and 0.1% NP-40) containing protease inhibitor cocktails (Nacalai Tesque). Supernatants obtained after centrifugation of the cell lysates were pre-cleared by incubation with protein A-Sepharose beads (GE Healthcare) at 4 °C for 30 min. After a brief centrifugation, supernatants were reacted at 4 °C overnight with Strep-Tactin sepharose beads (IBA Lifescience). The sepharose beads were collected by a brief centrifugation and washed once with high-salt buffer (1 M NaCl, 10 mM Tris-HCl [pH 8.0], 0.2% NP-40), twice with low-salt buffer (0.1 M NaCl, 10 mM Tris-HCl [pH 8.0], 0.2% NP-40), four times with radioimmunoprecipitation assay buffer (50 mM Tris-HCl [pH 7.5], 150 mM NaCl, 1% NP-40, 0.5% deoxycholate, 0.1% sodium dodecyl sulfate), and finally two times with UL13 kinase buffer (50 mM Tris-HCl [pH 8.0], 50 mM NaCl, 15 mM MgCl_2_, 0.1% Nonidet P-40, and 1 mM dithiothreitol). For *in vitro* kinase assays, UL13 kinase buffer containing 100 μM ATP and amylose beads containing purified MBP-EF-1δ (107–146) or MBP-EF-1δ (107–146)-S133A were added to the mixture of protein A-Sepharose beads and reacted at 30°C for 30 min. After incubation, the reaction mixture was mixed with 3x SDS sample buffer (187.5 mM Tris– HCl pH 6.5, 30% glycerol, 6% SDS, 15% 2-mercaptoethanol), boiled for 5 minutes, subjected to electrophoresis in denaturing gels. After electrophoresis, the separated proteins were transferred from the gels to nitrocellulose membranes (Bio-Rad), stained with Ponceau S, visualized on a ChemiDoc MP (Bio-Rad), and subjected to immunoblotting using the anti-Strep, anti-GFP or anti-EF-1δ-S133^P^ antibodies.

### Immunoprecipitation

Vero cells were infected with wild-type HSV-2 186 or YK873 (UL13-HA) at an MOI of 3 for 24 h and lysed in 0.1% NP40 buffer containing a protease inhibitor cocktail (Nacalai Tesque). After centrifugation, the supernatants were precleared by incubation with protein G-Sepharose beads, and then reacted with an anti-HA monoclonal antibody at 4°C for 2 h. Protein G-Sepharose beads were added to the supernatants, and the reaction continued for another 2 h. Immunoprecipitates were collected by a brief centrifugation, washed extensively with 0.1% NP-40 buffer, and analyzed by immunoblotting with the indicated antibodies.

### Inhibitor treatment

The proteasome inhibitor, MG132 (Wako), was added to the indicated COS-7 or HEK293T cells at 24 h after transfection at a final concentration of 10 μM.

### Determination of plaque size

Vero and U2OS cells were infected with each recombinant virus at an MOI of 0.0001, and plaque sizes were determined as described previously (30).

### Statistical analysis

Differences in viral replication and plaque size in cell cultures, and relative amounts of phosphorylated UL13 were analyzed statistically by analysis of variance (ANOVA) followed by Tukey’s post-hoc test. Differences in relative amounts of phosphorylated EF-1δ was evaluated by ANOVA followed by Tukey’s post-hoc test or one-way ANOVA with Dunnett’s multiple comparisons test comparing to the empty vector. A P value of <0.05 was considered statistically significant. All statistical analyses were performed with GraphPad Prism 8 (GraphPad Software, San Diego, CA).

## Acknowledgements

We thank Risa Abe, Tohru Ikegami, and Sachi Fujiwara for their excellent technical assistance. We are grateful to Yasuko Mori, Thomas Shenk, and Keiji Ueda for providing valuable reagents. This study was supported by Grants for Scientific Research and Grant-in-Aid for Scientific Research (S) (20H05692) from the Japan Society for the Promotion of Science, grants for Scientific Research on Innovative Areas (21H00338, 21H00417, 22H04803) and a grant for Transformative Research Areas (22H05584) from the Ministry of Education, Culture, Science, Sports and Technology of Japan, Precursory Research for Embryonic Science and Technology (JPMJPR22R5) from Japan Science and Technology Agency, grants (JP20wm0125002, JP22fk0108640, JP22gm1610008, JP223fa627001, JP23wm0225031, JP23wm0225035) from the Japan Agency for Medical Research and Development, grants from the International Joint Research Project of the Institute of Medical Science, the University of Tokyo, grants from the Takeda Science Foundation, the Mitsubishi Foundation, the Uehara Memorial Foundation, and the Waksman Foundation of Japan, and the GlaxoSmithKline Japan Research Grant 2019.

