## Supplementary material for "Identification of Viral Activators of the HSV-2 UL13 Protein Kinase": Table S1

**Table S1. Accession numbers of viral sequences using in this study.**

| Gene Simplex,Varicello,Mardi,ilto,Scuta | Accession Number | Link |
|---|---|---|
| Bovine alphaherpesvirus 5 (BoAHV5) | AY261359 | https://www.ncbi.nlm.nih.gov/nuccore/AY261359 |
| Bubaline alphaherpesvirus 1 (BuAHV1) | KU936049 | https://www.ncbi.nlm.nih.gov/nuccore/KU936049 |
| Bovine alphaherpesvirus 1 (IBRV) | JX898220 | https://www.ncbi.nlm.nih.gov/nuccore/JX898220 |
| Cervid alphaherpesvirus 1 (CvAHV1) | MH036942 | https://www.ncbi.nlm.nih.gov/nuccore/MH036942 |
| Cervid alphaherpesvirus 3 (CvAHV3) | MH036941 | https://www.ncbi.nlm.nih.gov/nuccore/MH036941 |
| Cervid alphaherpesvirus 2 (CvAHV2) | MH036943 | https://www.ncbi.nlm.nih.gov/nuccore/MH036943 |
| Caprine alphaherpesvirus 1 (CpAHV1) | MG989243 | https://www.ncbi.nlm.nih.gov/nuccore/MG989243 |
| Monodontid alphaherpesvirus 1 (MoAHV1) | MF678601 | https://www.ncbi.nlm.nih.gov/nuccore/MF678601 |
| Suid alphaherpesvirus 1 (PRV) | JF797218 | https://www.ncbi.nlm.nih.gov/nuccore/JF797218 |
| Equid alphaherpesvirus 8 (EqAHV8) | MF431611 | https://www.ncbi.nlm.nih.gov/nuccore/MF431611 |
| Equid alphaherpesvirus 9 (EqAHV9) | AP010838 | https://www.ncbi.nlm.nih.gov/nuccore/AP010838 |
| Equid alphaherpesvirus 1 (EAV) | AY665713 | https://www.ncbi.nlm.nih.gov/nuccore/AY665713 |
| Equid alphaherpesvirus 4 (EqAHV4) | AF030027 | https://www.ncbi.nlm.nih.gov/nuccore/AF030027 |
| Equid alphaherpesvirus 3 (EqAHV3) | KM051845 | https://www.ncbi.nlm.nih.gov/nuccore/KM051845 |
| Canid alphaherpesvirus 1 (CHV) | KT819633 | https://www.ncbi.nlm.nih.gov/nuccore/KT819633 |
| Phocid alphaherpesvirus 1 (PcAHV1) | MH509440 | https://www.ncbi.nlm.nih.gov/nuccore/MH509440 |
| Felid alphaherpesvirus 1 (FVRV) | FJ478159 | https://www.ncbi.nlm.nih.gov/nuccore/FJ478159 |
| Cercopithecine alphaherpesvirus 9 (SVV) | AF275348 | https://www.ncbi.nlm.nih.gov/nuccore/AF275348 |
| Human alphaherpesvirus 3 (VZV) | X04370 | https://www.ncbi.nlm.nih.gov/nuccore/X04370 |
| Gallid alphaherpesvirus 2 (MDV) | AF243438 | https://www.ncbi.nlm.nih.gov/nuccore/AF243438 |
| Gallid alphaherpesvirus 3 (GaAHV3) | HQ840738 | https://www.ncbi.nlm.nih.gov/nuccore/HQ840738 |
| Meleagrid alphaherpesvirus 1 (HVT) | AF291866 | https://www.ncbi.nlm.nih.gov/nuccore/AF291866 |

| | | |
|---|---|---|
| Columbid alphaherpesvirus 1 (PHV) | KX589235 | https://www.ncbi.nlm.nih.gov/nuccore/KX589235 |
| Anatid alphaherpesvirus 1 (DEV) | JF999965 | https://www.ncbi.nlm.nih.gov/nuccore/JF999965 |
| Spheniscid alphaherpesvirus 1 (SpAHV1) | LT608135 | https://www.ncbi.nlm.nih.gov/nuccore/LT608135 |
| Cercopithecine alphaherpesvirus 2 (SA8) | AY714813 | https://www.ncbi.nlm.nih.gov/nuccore/AY714813 |
| Papiine alphaherpesvirus 2 (HPV2) | DQ149153 | https://www.ncbi.nlm.nih.gov/nuccore/DQ149153 |
| Macacine alphaherpesvirus 1 (BV) | AF533768 | https://www.ncbi.nlm.nih.gov/nuccore/AF533768 |
| Human alphaherpesvirus 2 (HSV2) | JN561323 | https://www.ncbi.nlm.nih.gov/nuccore/JN561323 |
| Panine alphaherpesvirus 3 (ChHV) | JQ360576 | https://www.ncbi.nlm.nih.gov/nuccore/JQ360576 |
| Human alphaherpesvirus 1 (HSV1) | JN555585 | https://www.ncbi.nlm.nih.gov/nuccore/JN555585 |
| Teropodid alphaherpesvirus 1 (FBAHV1) | AB825953 | https://www.ncbi.nlm.nih.gov/nuccore/AB825953 |
| Macropodid alphaherpesvirus 1 (MaAHV1) | KT594769 | https://www.ncbi.nlm.nih.gov/nuccore/KT594769 |
| Macropodid alphaherpesvirus 2 (MaAHV2) | MT900475 | https://www.ncbi.nlm.nih.gov/nuccore/MT900475 |
| Bovine alphaherpesvirus 2 (BMV) | MT862163 | https://www.ncbi.nlm.nih.gov/nuccore/MT862163 |
| Leporid alphaherpesvirus 4 (LHV4) | JQ596859 | https://www.ncbi.nlm.nih.gov/nuccore/JQ596859 |
| Ateline alphaherpesvirus 1 (HVA1) | KY385637 | https://www.ncbi.nlm.nih.gov/nuccore/KY385637 |
| Saimiriine alphaherpesvirus 1 (HVS1) | HM625781 | https://www.ncbi.nlm.nih.gov/nuccore/HM625781 |
| Gallid alphaherpesvirus 1 (ILTV) | JN596962 | https://www.ncbi.nlm.nih.gov/nuccore/JN596962 |
| Psittacid alphaherpesvirus 1 (PDV) | AY372243 | https://www.ncbi.nlm.nih.gov/nuccore/AY372243 |
| Chelonid alphaherpesvirus 5(FPTHV) | HQ878327 | https://www.ncbi.nlm.nih.gov/nuccore/HQ878327 |
| Testudinid alphaherpesvirus 3(TeHV3) | KM924292 | https://www.ncbi.nlm.nih.gov/nuccore/KM924292 |
